# Deep Learning-Enhanced 3D Imaging Unveils Semaglutide Impact on Cardiac Fibrosis

**DOI:** 10.1101/2025.02.25.640039

**Authors:** Sheyla Barrado-Ballestero, Sarah Torp Yttergren, Max Hahn, Marie Biviano Rosenkilde, Ditte Marie Jensen, Michael Christensen, Louise Thisted, Heidi Lindgreen Holmberg, Geoffrey Teixeira, Tor Biering-Sørensen, Casper Gravesen Salinas, Urmas Roostalu

**Affiliations:** Gubra ApS; Hørsholm Kongevej 11B, 2970 Hørsholm, Denmark; Department of Biomedical Sciences, University of Copenhagen; Blegdamsvej 3B, 2200, Denmark; Alentis Therapeutics; Hegenheimermattweg 167A, 4123 Allschwil, Switzerland; Center for Translational Cardiology and Pragmatic Randomized Trials, Department of Biomedical Sciences, University of Copenhagen; Blegdamsvej 3B, 2200, Denmark; Cardiovascular Non-Invasive Imaging Research Laboratory, Copenhagen University Hospital - Herlev and Gentofte; Steno Diabetes Center Copenhagen; Department of Cardiology, Copenhagen University Hospital – Rigshospitalet

## Abstract

**Background:** Extensive preclinical research aims to develop novel therapeutics for myocardial fibrosis (MF), a condition marked by collagen accumulation that impairs cardiac function. MF is particularly relevant in heart failure with preserved ejection fraction (HFpEF), a growing clinical challenge with limited treatment options. However, current methods for quantifying MF in mouse models struggle to accurately capture its heterogeneous regional distribution, creating a significant barrier to reliably assessing the efficacy of therapeutics.

**Purpose:** To develop a whole-heart fibrosis imaging and deep learning (DL)-based quantification method and validate the workflow by assessing the efficacy of a glucagon-like peptide-1 receptor (GLP-1R) agonist in mouse HFpEF model.

**Experimental Approach:** By utilizing a fluorescent collagen-labelling dye, tissue clearing and 3D light sheet microscopy, we developed a high-throughput imaging platform for MF. We established DL framework to quantify perivascular and replacement fibrosis, as well as hypertrophy, in 17 left ventricular (LV) segments. The antifibrotic effects of the GLP-1R agonist semaglutide were evaluated in the db/db UNx-ReninAAV mouse model, which exhibits diabetes, kidney failure, obesity, and hypertension.

**Key Results:** Whole-heart 3D light sheet microscopy, combined with artificial intelligence, enables micrometer-resolution analysis of MF distribution in rodents. This approach allows for detailed characterization of distinct regional fibrosis patterns. Chronic semaglutide treatment significantly reduced LV hypertrophy and perivascular fibrosis but had no significant effect on replacement fibrosis.

**Conclusions and Implications:** The established 3D imaging and quantification approach provides a powerful tool for evaluating the therapeutic efficacy of antifibrotic compounds and studying the cellular and pathological mechanisms underlying cardiovascular diseases.

## Introduction

Heart failure (HF) is the leading cause of death worldwide. Despite advances in the development of therapeutics, morbidity and mortality rates remain high. Notably, the incidence of HF with preserved ejection fraction (HFpEF) has been increasing, now accounting for approximately half of all HF cases (Clark et al., 2022). HFpEF is a heterogeneous condition that develops over a prolonged period and is driven by diverse risk factors, including chronic and poorly controlled hypertension, obesity, type 2 diabetes, and advanced age (Savji et al., 2018; Martinez-Morata et al., 2024). While the pathophysiological mechanisms of HFpEF are not fully understood, myocardial fibrosis (MF) is consistently recognized as a hallmark feature, strongly correlating with disease severity and outcomes (Kato et al., 2015a; Schelbert et al., 2017; Kanagala et al., 2019; López et al., 2021). Fibrosis in HFpEF often precedes overt HF symptoms, progressing gradually over time. In its early stages, MF typically manifests as diffuse extracellular matrix accumulation between myofibers and around coronary vasculature (perivascular fibrosis). As the disease progresses, these fibrotic regions expand, increasing tissue stiffness and impairing normal cardiac contraction. Advanced fibrosis leads to cardiac myocyte necrosis and subsequent replacement fibrosis in necrotic areas, a process largely considered irreversible (Mewton et al., 2011; Díez et al., 2020; Sweeney et al., 2020). While HFpEF risk factors can independently contribute to HF and MF, the interplay of multiple comorbidities and their impact on fibrosis progression and treatment efficacy remain poorly understood (Mall et al., 1990; van Hoeven and Factor, 1990; Kawaguchi et al., 1997; Izumaru et al., 2019; Pop-Busui et al., 2022; González et al., 2024). Notably, no pharmacological agents specifically targeting MF have yet reached clinical application.

Quantitative assessment of MF remains a significant challenge in both clinical and preclinical research, hindered by technical limitations and the inherent variability of the disease (Barton et al., 2023; Bengel et al., 2023; Karur et al., 2024). MF analysis in preclinical rodent models is complicated by the small size of the heart and lack of methods that provide sufficient resolution (Galati et al., 2016). Histological staining techniques, such as Masson’s Trichrome staining, which relies on Fast Green (FG) dye to label collagen, remain the gold standard for MF assessment. However, due to the heterogeneous distribution of MF, analyzing one or a few histological sections from the heart often fails to accurately capture the extent of fibrosis or to reliably quantify the effects of novel therapeutics and describe disease mechanisms. Light sheet fluorescence microscopy (LSFM) has recently emerged as a powerful tool for imaging entire rodent hearts at single-cell resolution (Fei et al., 2016; Ding et al., 2017; Merz et al., 2019; Roostalu et al., 2021), but the method has not been applied in fibrosis analysis, due to technical limitations of achieving uniform tissue labelling and lack of image analysis tools. FG, a small dye that fluoresces in the far-red wavelength (Timin and Milinkovitch, 2023), presents a promising candidate for developing a 3D MF imaging method.

Semaglutide, a glucagon-like peptide-1 (GLP-1) receptor agonist is an efficacious treatment option for diabetes and obesity (Wilding et al., 2021; Lincoff et al., 2023). Accumulating evidence supports the beneficial effects of semaglutide in the management of cardiovascular diseases. Semaglutide reduces HF symptoms in obesity-associated HFpEF and diminishes major adverse cardiovascular event (MACE) risks and mortality in both HFpEF and HFrEF (Kosiborod et al., 2023, 2024; Deanfield et al., 2024). To what extent these beneficial clinical effects encompass MF reduction has not been studied. In a mouse model of HFpEF, resulting from obesity and hypertension (infusion of angiotensin II, AngII), GLP-1 receptor agonists were found to reduce MF and hypertrophy (Withaar et al., 2021, 2023; He et al., 2024). The *db/db* UNx-ReninAAV mouse model, developing kidney failure, diabetes, obesity and hypertension, has gained popularity in assessing cardiorenal endpoints in drug discovery research (Harlan et al., 2018; Østergaard et al., 2021a; Dalbøge et al., 2022). We hypothesized that the gradual progression of the disease in this mouse model that combines all the key risk factors of HFpEF makes it relevant to study the effect of semaglutide in reducing MF.

Here, we present a high-throughput LSFM and artificial intelligence (AI) image analysis platform for the quantitative assessment of cardiac hypertrophy and fibrosis in entire rodent hearts. Using a rapid staining protocol with FG, collagen is visualized and quantified in three-dimensional (3D) space throughout whole hearts. The analysis differentiates between interstitial, replacement, and perivascular fibrosis, providing comprehensive insights into fibrosis subtypes. To enhance spatial specificity, we adapted the 17-region anatomical model from the American Heart Association (AHA) nomenclature for use in light sheet-imaged mouse hearts. This adaptation enables detailed quantification of regional variations in MF. Applying this platform to the db/db UNx-ReninAAV mouse model, we demonstrated that semaglutide treatment effectively reduced cardiac hypertrophy and mitigated the progression of perivascular fibrosis.

## Materials & Methods

### Ethics statement and Animals

The Danish Animal Experiments Inspectorate approved all experiments, which were conducted using internationally accepted principles for the use of laboratory animals (License No. 2023-15-0201-01456). Female db/+ and db/db (BKS.CgDock7mþ /þ Leprdb/J) mice (5-wk-old at arrival and 12-week-old at study start) were obtained from Charles River (Calco, Italy) and housed in a controlled environment (12:12-h light-dark cycle, 23 ± 2C, humidity: 50 ± 10%). Each animal was identified by an implantable subcutaneous microchip (PetID Microchip, E-vet, Haderslev, Denmark). Mice had ad libitum access to standard chow (Altromin 1324, Brogaarden, Hørsholm, Denmark) and tap water.

### In vivo pharmacology

Female db/+ mice received a single intravenous dose of ReninAAV (1.5×10^10^ GC) by study week -5 and underwent unilateral nephrectomy (UNx) at study week -4, resulting in diabetic nephropathy hypertensive mice (DN/HT) (Østergaard et al., 2021b; Dalbøge et al., 2022). A group of Healthy controls (female db/+) was included in the study (n=9). Prior to treatment, DN/HT mice were randomized based on body weight, fed blood glucose (BG) and urinary albumin-to-creatinine ratio (uACR) to Vehicle (n=18) and Semaglutide (n=16) treatment arms. Semaglutide (Ozempic, Novo Nordisk) was administered subcutaneously (SC) daily for 12 weeks, and dosing was gradually up-titrated the first days of treatment: 3 nmol/kg (day 1-3), 10 nmol/kg (day 4-6) and 20 nmol/kg (from day 7). The Healthy control and DN/HT vehicle groups were dosed SC daily with vehicle. Terminal endpoints included blood HbA1c and plasma urea. The heart was weighed, and further processed for 3D imaging. Tibial length was determined for normalization of heart weight.

### Plasma and urine biomarker sampling and analysis

For measurement of blood glucose, blood was collected by tail vein puncture, stored in heparinized glass capillary tubes and submerged in 500 µl glucose/lactate system solution buffer (EKF-diagnostics, Germany). Blood glucose was measured using a BIOSEN c-Line glucose meter (EKF-diagnostics, Germany) according to the manufacturer’s instructions. For the HbA1c analysis, blood samples were collected by tail vein puncture on the morning of each termination, placed into heparinized glass capillary tubes and immediately suspended in Hemolyzing Reagent (Roche Diagnostics) and stored at -70°C until analysis. HbA1c was measured using commercial kit (Roche Diagnostics,) on the Cobas c 501 autoanalyzer. Urine was collected from a clean cage in 0.5 ml Eppendorf tubes and stored at -70°C until analysis. Samples were centrifuged at 2000 g for 2 min prior to analysis. Albumin was measured using a commercial ELISA kit (Bethyl Laboratories, Inc.). Creatinine was measured using a commercial kit (Roche Diagnostics) on the Cobas c 501 autoanalyzer.

### Tissue extraction and preparation for LSFM

For 3D heart imaging, the heart was collected immediately after terminal blood sampling. Following excision, the heart was perfused via the aorta and vena cava with freshly prepared cardioplegic solution (St. Thomas’ Hospital cardioplegic solution no. 2: 110.0 mM NaCl, 10.0 mM NaHCO3, 16.0 mM KCl, 16.0 mM MgCl2, 1.2 mM CaCl2; pH 7.8) to thoroughly flush out blood from the cardiac chambers. The heart was then immersed in the same cardioplegic solution for approximately 30 seconds, inducing contraction. After blotting dry, the heart was weighed and fixed in 10% neutral buffered formalin and stored at room temperature. The kidney and liver are dissected and quickly washed in PBS to remove excessive blood and fixed overnight at room temperature.

### Collagen staining in whole mouse organs

After overnight fixation, the heart, kidney and liver were washed in phosphate buffered saline (PBS) for 1 hour and dehydrated through a methanol/H2O gradient (20%, 40%, 60%, 80% and 100% methanol (VWR International A/S), each for 1 hour at room temperature. The next day, they were washed twice in 100% methanol for 30 minutes, cooled to 4°C for 1 hour, and bleached overnight in freshly prepared 5% H_2_O_2_ (Acros Organics, Fisher Scientific Biotech Line A/S, Slangerup, Denmark) in methanol (1 part 35% H2O2 to 6 parts methanol) at 4°C. This bleaching step was repeated once more overnight at 4°C. On the following day, the samples were washed twice in 100% methanol for 1 hour and incubated for 6 days in 0.1 mg/ml Masson’s trichrome (Fast Green FCF, Thermo Fisher Scientific, CAS 2353-45-9, cat. A16520.06) in methanol. After staining, samples were washed in 100% methanol for 1 hour, followed by two washes of 2 hours each, and then left overnight at room temperature. All steps were conducted in tightly sealed tubes to prevent evaporation and oxidation. For tissue clearing, samples were incubated overnight in 100% methanol, followed by 3 hours in 66% Dichlormethane (DCM, Sigma-Aldrich, CAS 75-09-2)/33% methanol at room temperature with orbital shaking. They were then treated in 100% DCM for two 1-hour intervals to ensure methanol removal, before being transferred to ethyl cinnamate (ECi) for storage in dark, sealed glass vials.

### Light-sheet fluorescence microscopy of cleared tissues and post-processing

For quantitaive light sheet imaging, heart samples were imaged using a Bruker Luxendo LCS SPIM equipped with a Hamamatsu ORCA-Flash4.0 V3 digital sCMOS camera and with ECi as the clearing agent during data acquisition. Images were acquired at dual illumination excitation with 4.4× magnification (rendering 1.4625µm^2^ per pixel) and z-step size of 10µm. FG collagen signal was captured at 642nm wavelength for excitation and bandpass emission filter between 655-704nm with an exposure time of 45ms, beam expander of 6.3µm and tunable acoustic gradient (TAG) lens at 35% to homogenize uniformly the light-sheet across the FOV. The natural occurring autofluorescence signal to outline the cardiac anatomy was captured at 561nm excitation and a bandpass 580-627nm emission filter with an exposure time of 40ms, beam expander of 6.3µm and TAG lens at 35%. To capture the entire heart morphology, 4 tiles were used along the X-dimension and 5 tiles along the Y-dimension, resulting in an average of 686 slices in the Z-dimension.

For high resolution scans of selected regions, a LaVision ultramicroscope II setup (Miltenyi Biotec, Bergisch Gladbach, Germany) equipped with a pco.edge 4.2 sCMOS camera (PCO imaging AG, Kelheim, Germany), SuperK EXTREME supercontinuum white-light laser EXR-15 (NKT Photonics, Birkerød, Denmark) and MV PLAPO 2XC (Olympus, Tokyo, Japan) objective lens was used. Hearts were mounted to the sample holder using clear 514 silicone (Dana Lim, Køge, Denmark) and imaged in ECi. Version 7 of the Imspector microscope controller software was used to acquire images at 6.3× magnification (12.6× total magnification), with an exposure time of 29.6 ms for FG collagen (630nm excitation and 680nm emission) and the same exposure time for autofocus channel (500nm excitation and 545nm emission), in a z-stack with 3-µm intervals and sheet thickness of 3.43µm, resulting a final size of 0.48×0.48×3µm per voxel.

### AI-Quantification of cardiac fibrosis

To accurately quantify fibrosis and cardiac morphometry in whole rodent hearts, we developed a two-stage quantification. The first stage focuses on large-scale segmentation and alignment of the heart tissue at low resolution. The second stage provides a detailed analysis of collagen distribution at high resolution. Finally, we merged the outputs of both pipelines to achieve high-resolution regional quantification of fibrosis throughout the heart.

In both pipelines, we used an end-to-end deep learning model based on a residual U-Net (ResUNet) architecture, implemented within the TensorFlow-n framework (Abadi et al., 2016). The architecture consists of an input layer matching the input dimensions, followed by an encoder path composed of four down sampling stages. Each stage includes a residual block with filters set to N, 2N, 4N, and 8N (where N is the base number of filters). These residual blocks, utilizing a kernel size of 3×3×3, are designed to enhance feature extraction while preserving essential spatial details through residual connections. After each residual block, down sampling is applied via a max-pooling layer with a window size of 2×2×2. The bottleneck layer, positioned at the deepest level of the network, utilizes a residual block with 16×N filters to capture abstract, high-level features necessary for segmentation. In the decoder path, the up sampling begins with the bottleneck layer output. Each up sampling stage uses a layer of (2×2×2) followed by concatenation with the corresponding encoder layer, maintaining high-resolution feature details. Each concatenation is followed by a residual block with progressively reduced filter sizes (8×N, 4×N, 2×N, N) to recover spatial resolution progressively. A batch normalization layer is included after each pooling operation in the encoder to stabilize training. The output layer applies a 1×1×1 convolution with softmax activation to generate the class probabilities for each pixel or voxel. All the training parameters were established using two key callbacks that monitor training progress: *ReduceLROnPlateau*, which reduces the learning rate by a factor of 0.1 after 10 epochs of plateaued validation loss, and *EarlyStopping*, which halts training after 20 epochs without improvement, restoring the best weights based on the lowest validation loss.

The first step in the low-resolution pipeline consisted of preprocessing the autofluorescence channel (561 nm) for heart region segmentation. Since the pixel intensities across the heart are uniform and there is no physical distinction between the different regions, the whole sample was required as model input. We first created a bounding box around the tissue to mask out background regions. We resampled the voxel size to isotropic resolution to ensure uniform scaling in all directions. Then, we resample the isotropic volume to the desired input size [128, 128, 128], ensuring the entire volume fits within the model. Final steps involve applying a square root transformation and rescaling the volume intensities to the 0-1 range.

We then proceeded to manual data annotation of the regions of interest (LV wall and chamber) using the ITK-SNAP active contour segmentation tool. Using the ResUNet from above, we trained and validated the segmentation model using a dataset of 26 samples, achieving an accuracy of 85%. The model targets three classes: LV wall, LV chamber, and background. It incorporates N=16 filters and a batch size of 4 samples. We used Volumentations to augment the training data (Solovyev et al., 2022), via randomly generated transforms, including flips, brightness/contrast adjustments, and Gaussian blur to enhance generalization. The model was trained using Dice loss and Adam optimizer with a learning rate of 2e^-4^ over 300 epochs.

After training, we predicted the remaining hearts and proceeded to the extraction of the 17 segments from the LV wall. Our procedure starts with the geometrical alignment of the hearts to a common orientation. We determined the primary axis of the heart by applying a principal component analysis (PCA) based on the distribution of vertices of the surface geometry obtained using marching cubes (Lewiner et al., 2003). The heart was rotated in two stages to align the apex in the inferior area of the volume, followed by a final rotation along the short axis to ensure the LV was positioned on the left side.

Finally, the LV mask was divided into 17 regions based on the AHA segmentation model. The voxel coordinates of all non-zero points in the LV mask are used to compute the range of the projection along the principal axis. The mask is then divided into three regions—basal, mid, and apical—using this projection. The basal, mid, and apical regions are further subdivided into 6, 6, and 4 segments, respectively (with an additional apex segment). The voxel coordinates are transformed into polar coordinates and labels are assigned to each segment accordingly.

Similarly, in the high-resolution pipeline, the specific channel (642 nm) was processed for high-resolution regional quantification of fibrosis throughout the heart. First, we divided the input volumes into smaller patches and extracted the middle slice of each. Next, we generated embeddings using OpenAI PLIP for data exploration based on the clean vision scores (blurriness, darkness, and low information). After filtering, we annotated the selected patches using Fiji LabKit plugin from (Arzt et al., 2022). For model training, each volume was subdivided into smaller patches of [64, 96, 96] voxels and preprocessed with square root transformation to improve model accuracy and stability, and then rescaled to the unit range [0, 1]. Same as before, we included an augmentation step to reduce overfitting on the training dataset and improve generalizability on the testing dataset. We used 80% of the patches for training and 20% for validation using the ResNet model with the same configuration as described earlier except for using a batch size of 16 and dice plus focal loss. Moreover, to deal with the imbalanced dataset, we gave extra weight to the minority class (fibrosis) during the loss computation. We repeated the same procedure but annotated all the collagen fibers to segment the whole collagen network.

As the last step, fibrosis was measured in cardiac segments. We upsampled the label map with the 17 segments and registered it to the specific channel using rigid transform from ITK-Elastix (Ntatsis et al., 2023). In the LV mask, we applied a binary erosion and multiplied it to the fibrosis segmentation to remove the heart membrane. In the prediction, we removed small non-connected structures, we filled the blood vessels and computed the local thickness to split the perivascular fibrosis. Once cleaned, we measured the regional fibrosis content as the number of non-zero pixels from the predicted fibrosis in that region divided by the region size. We also measured collagen density, and whole fibrosis content *(mm*^3^). All statistical analyses were performed in GraphPad Prism (version 8.4.2). Dunnett test was used for statistical comparison against the *DN/HT* mouse model with vehicle after ANOVA. All data are presented as mean values and standard errors of the mean (s.e.m).

## Results

### Model characterization study

This study aimed to develop a rapid cardiac fibrosis analysis method for drug discovery and apply it to analyze the impact of semaglutide on MF. First, we sought suitable mouse models that carry the common co-morbidities of HFpEF and develop MF (Figure 1A). In a pilot experiment, we conducted a time-series analysis of the *db/db* UNx-ReninAAV model, from now on DN/HT (diabetic nephropathy and hypertension), describing key cardiometabolic parameters at day 1, and 4, 8 and 12 weeks following UNx surgery, compared to *db/+* control group, from now on Healthy control (age-matched to the 12-week study group). At all analyzed time points the body weight in the DN/HT model was higher than in the Healthy control group, with no significant differences between the time points (Figure 2A). An increase in HbA1c was evident from 4 weeks after model induction (p<0.001) indicating development of diabetes (Figure 2B). Heart weight was also significantly increased after week 4 (Figure 2C). Kidney failure progression was monitored by urine albumin-to-creatine ratio (ACR). The DN/HT mice showed increased ACR at all analyzed time points with no significant differences between them (p<0.0001 compared to Control; Figure 2D). Altogether these data indicate established diabetes, kidney failure, cardiac hypertrophy and obesity at 4 weeks, with stable disease parameters from 8 to 12 weeks. Analyzing the 12-week time-point allows thus an 8-week treatment window in the DN/HT model with established underlying pathology for studying the efficacy of pharmaceuticals to prevent and reverse disease endpoints.

**Figure 1.**
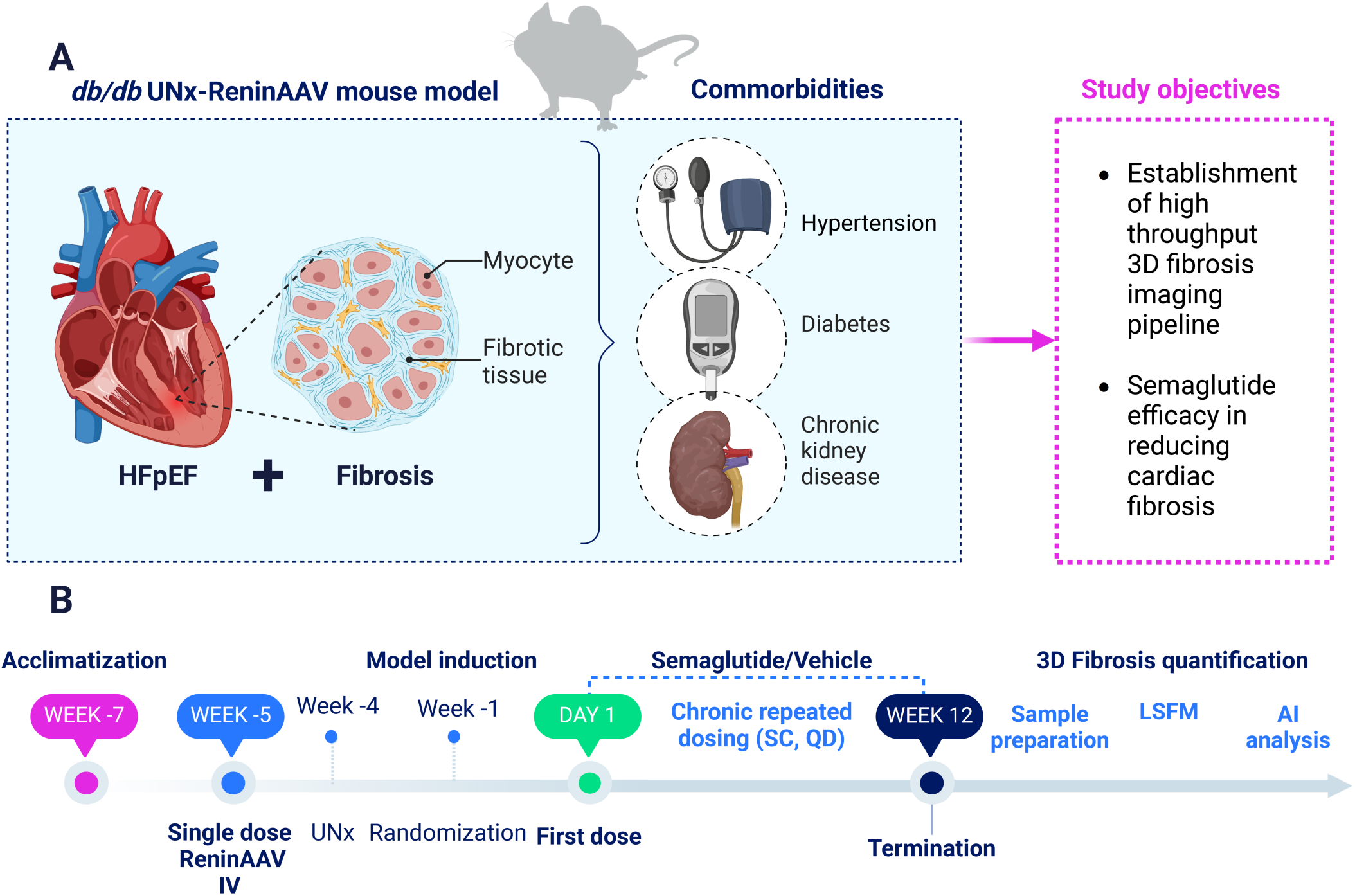
(A) Establishment of a mouse model with hypertension, kidney failure, obesity and diabetes, leading to HFpEF and cardiac fibrosis. (B) Study outline overview to characterize the effect of semaglutide on the *db/db* UNx-ReninAAV (DN/HT) mouse model. Abbreviations: IV – intravenous, LSFM: light sheet fluorescence microscopy, QD – once daily, SC – subcutaneous, UNx – uninephrectomy. Schematic image was generated with BioRender.

**Figure 2.**
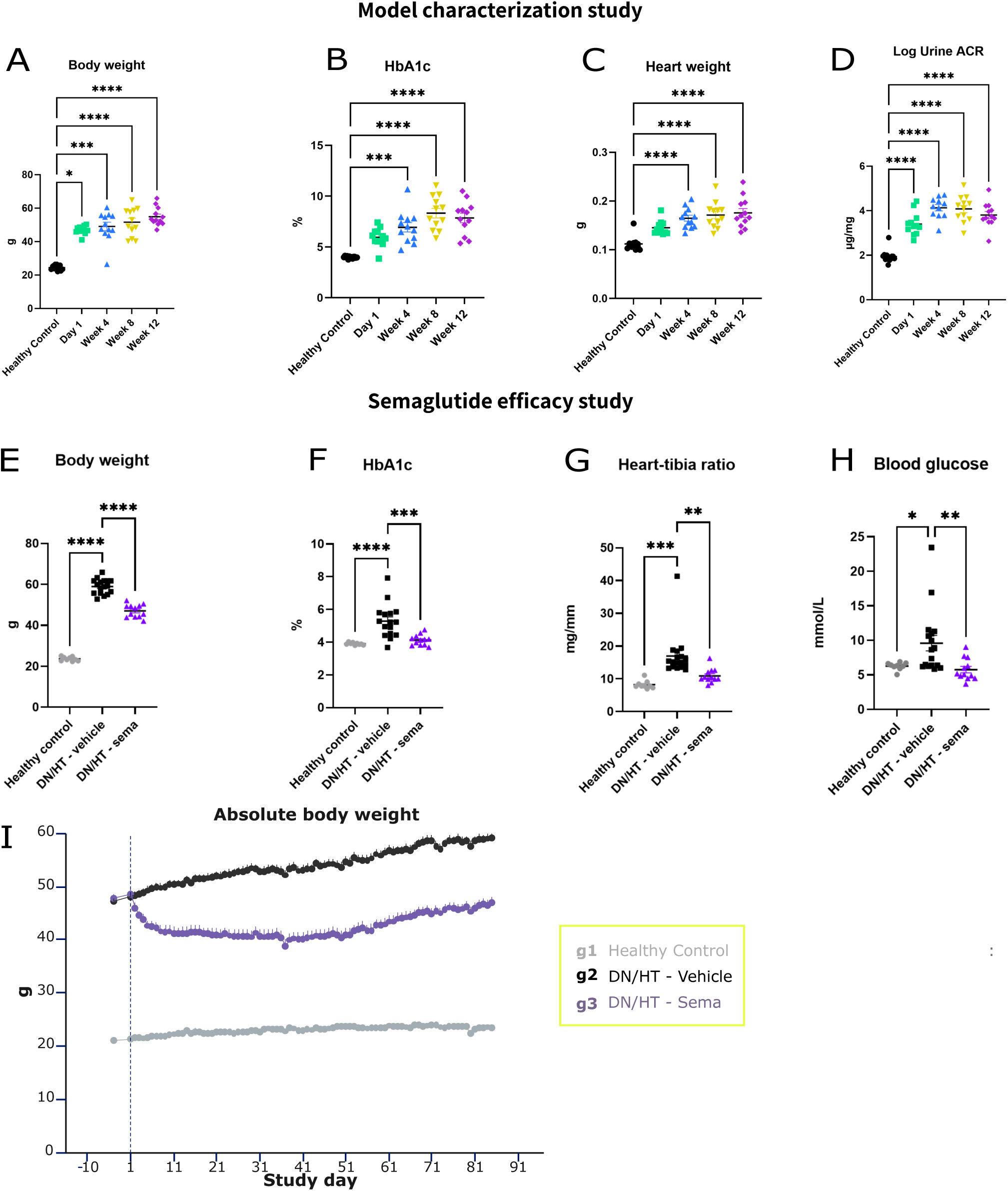
Pharmacokinetics for model characterization and Semaglutide efficacy. (A-I). Establishment of obesity, diabetes and kidney disease in the *db/db* UNx-ReninAAV (DN/HT) mouse model (A-C). (A) Body weight (BW) measured at termination. (B) Hemoglobin A1C (HbA1c) data for last study day. (C) Urine albumin to creatinine ratio (Urine ACR) at study end. Control: n= 13, Week (Wk) 4 to Wk 16: n= 12 for BW (A), HbA1c (B) and Urine ACR (C), except Wk 8 Urine ACR n=11. Kruskal-Wallis test with Dunn’s test for multiple comparisons was used for BW (A) and HbA1c (B) data. One-way ANOVA with Dunnet’s post-hoc test to compare means of each group to the control group was applied to log-transformed Urine ACR (C) data. Cardiometabolic effects of vehicle and semaglutide treatment (n=12-17) (G-K). (G) BW was measured every day (J) and at (K) termination. (L) HbA1c, (J) Heart-tibia ratio and (K) blood glucose levels were determined at last study day. Data is presented as mean ± SEM. One-way ANOVA with Dunnet’s post-hoc test to compare means of each group to db/db UNx-ReninAAV Vehicle. *p<0.05; ***p<0.001; ****p<0.0001 vs Control. Graphs are generated using GraphPad Prism (v 9.2).

### Semaglutide efficacy study

By considering the above defined characteristics of the DN/HT mouse model, we next proceeded with a semaglutide efficacy study where treatment of MF was the primary endpoint (Figure 1B). Semaglutide (DN/HT – sema) and vehicle (DN/HT – vehicle) were dosed daily over the 12-week study period. Age-matched *db/+* mice served as a control group (Healthy control). The final sample size included in 3D imaging was n=9 for Healthy control, n=17 DN/HT-vehicle and n=12 DN/HT-sema. As secondary endpoints, we assessed key metabolic parameters. On average, the Healthy control group gained 10.5% and DN/HT group 23.0% in body weight over the treatment period, whereas semaglutide treatment resulted in 3% loss in body weight compared to study start (Figure 2I). The final body weight at termination was 47±3.2 g in DN/HT – sema, compared to 59±3.6 g in DN/HT – vehicle (p<0.001) and 23.6±1.01 g in the Healthy control (Figure 2E). Blood HbA1c (%) analysis at study end demonstrated a reduction from 5.3 +/- 1.03 in the DN/HT – vehicle group to 4.1 +/- 0.3 in the DN/HT – sema (p<0.001), which was similar to the Healthy control mean value (3.9±0.1) (Figure 2F). Normalized heart weight was significantly decreased in the semaglutide treated group in comparison to the vehicle treated group (DN/HT – sema: 10.9±2.2; DN/HT – vehicle: 17±6.6; Healthy control: 8.2 ±1.2) (Figure 2G). Similarly, blood glucose levels in the DN/HT- Sema group at study end were significantly lower in comparison to DN/HT – vehicle, and comparable to the level in the Healthy control group (6.3±0.2 mmol/l in Healthy control; 9.6±1.1 mmol/l in DN/HT – vehicle; 5.8±0.5 mmol/l in DN/HT-Sema) (Figure 2H).

### Rapid 3D imaging and AI-Quantification of cardiac fibrosis

We developed a method for visualizing collagen accumulation with micrometer-level resolution in whole organs using FG staining, tissue clearing and LSFM. Whole mouse hearts were made optically transparent using ethyl cinnamate (ECi) to match the refractive index of fixed proteins and lipids (Figure S1). A 3D whole-heart reconstruction of a Healthy control mouse (Figure 3A-E) reveals the collagen network supporting the heart’s structure within the myocardium. Chordae tendinea demonstrated dense collagen fibers, whereas thin fibers run in parallel to cardiomyocytes (Figure 3B-C). A dense collagen network encases the outer surface of the heart and major blood vessels (Figure 3D-E). As proof of concept, the collagen imaging method was applied to the entire mouse liver and kidney, supporting broader applicability beyond cardiac analysis (Figure S2 and S3; Video 1 and 2). Application of the method on hearts from the DN/HT – vehicle group revealed replacement as well as perivascular fibrosis (Figure 3F-H; M). The collagen network in-between cardiomyocytes was qualitatively similar in all study groups. Replacement fibrosis could also be detected in the hearts from DN/HT – sema group (Figure 3I-K). Qualitative assessment suggested a reduction of perivascular fibrosis in the DN/HT – sema group in comparison to DN/HT – vehicle, (Figure 3L-N, Video 3).

**Figure 3.**
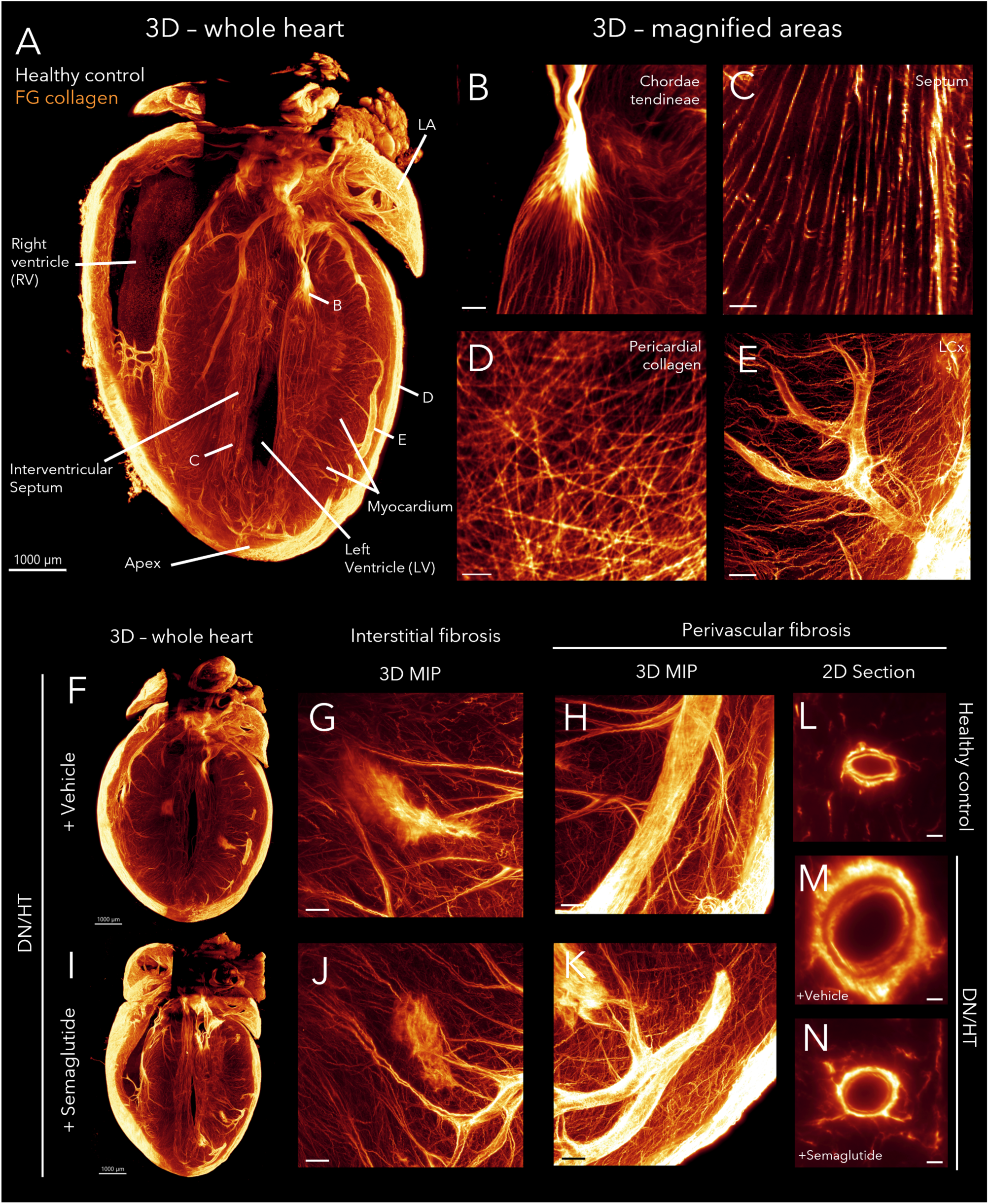
3D light sheet fluorescence microscopy reveals cardiac fibrosis and the effects of semaglutide treatment in diabetic mouse hearts with Fast Green (FG) collagen staining. (A-E) Representative 3D light sheet fluorescence microscopy (LSFM) images of collagen distribution in a healthy Control (*db/+)* mouse heart, labeled using FG to visualize collagen fibers in glow color. (A) A maximum intensity projection (MIP) of a 1 mm thick slice through the 3D imaged heart in long-axis plane, shows the left ventricle (LV), right ventricle (RV), myocardium, interventricular septum, apex left atrial appendage (LA) and chordae tendineae, with FG highlighting collagen. (B-E) Magnified views display specific collagen-rich structures: chordae tendineae (B), collagen fibers between cardiomyocytes of the interventricular septum (C), pericardial collagen network (D), and perivascular collagen of the left circumflex artery (LCx) (E). (F-K) Representative LSFM images of *db/db* UNx-ReninAAV (DN/HT) whole mouse hearts treated with vehicle (F-H) or semaglutide (I-K) reveal increased collagen deposition, evidenced by increased tissue volume and enhanced interstitial and replacement fibrosis (G and J)) and perivascular fibrosis (H and K), both visualized using 3D MIPs. (L-N) 2D cross-sectional images of perivascular collagen surrounding a LCx in *db/+* (L), *db/db* UNx-ReninAAV + Vehicle (M), and *db/db* UNx-ReninAAV + Semaglutide (N) hearts. Scale bars are 1000µm (A, F, I), 100µm in (B-E, G-H, J-K) and 20µm in (L-N).

We next proceeded to develop a quantification workflow for cardiac fibrosis analysis (Figure 4). First, we sought to implement a pipeline to obtain cardiac morphometrical endpoints (LV mass [mm3], LV chamber volume [mm3] and local thickness [mm] (Dahl and Dahl, 2023). After preprocessing the autofluorescence (AF) channel, we implemented a deep learning (DL)-based framework to segment the LV wall and chamber using a 3D U-Net for pixel semantic segmentation. As qualitative assessment indicated large regional variability, we sought to further divide the LV wall into sub-regions. To keep consistency with the human cardiac nomenclature and clinical data, we subdivided the LV according to the segmentation proposed by the American Heart Association (AHA) (Cerqueira et al., 2002) into three sections: base, mid-cavity, and apex. In the following step, each region was further subdivided to obtain the 17 segments: 6 basal segments, 6 mid-cavity segments, 4 apical segments, and the true apex as segment 17. In parallel, we predicted separate masks from the specific FG channel to segment the total collagen content, perivascular fibrosis and replacement fibrosis. The deep learning identification of collagen network helped to characterize the natural interstitial pattern seen in the myocardium. The stringent identification of fibrosis helped to identify both interstitial and perivascular fibrosis.

**Figure 4.**
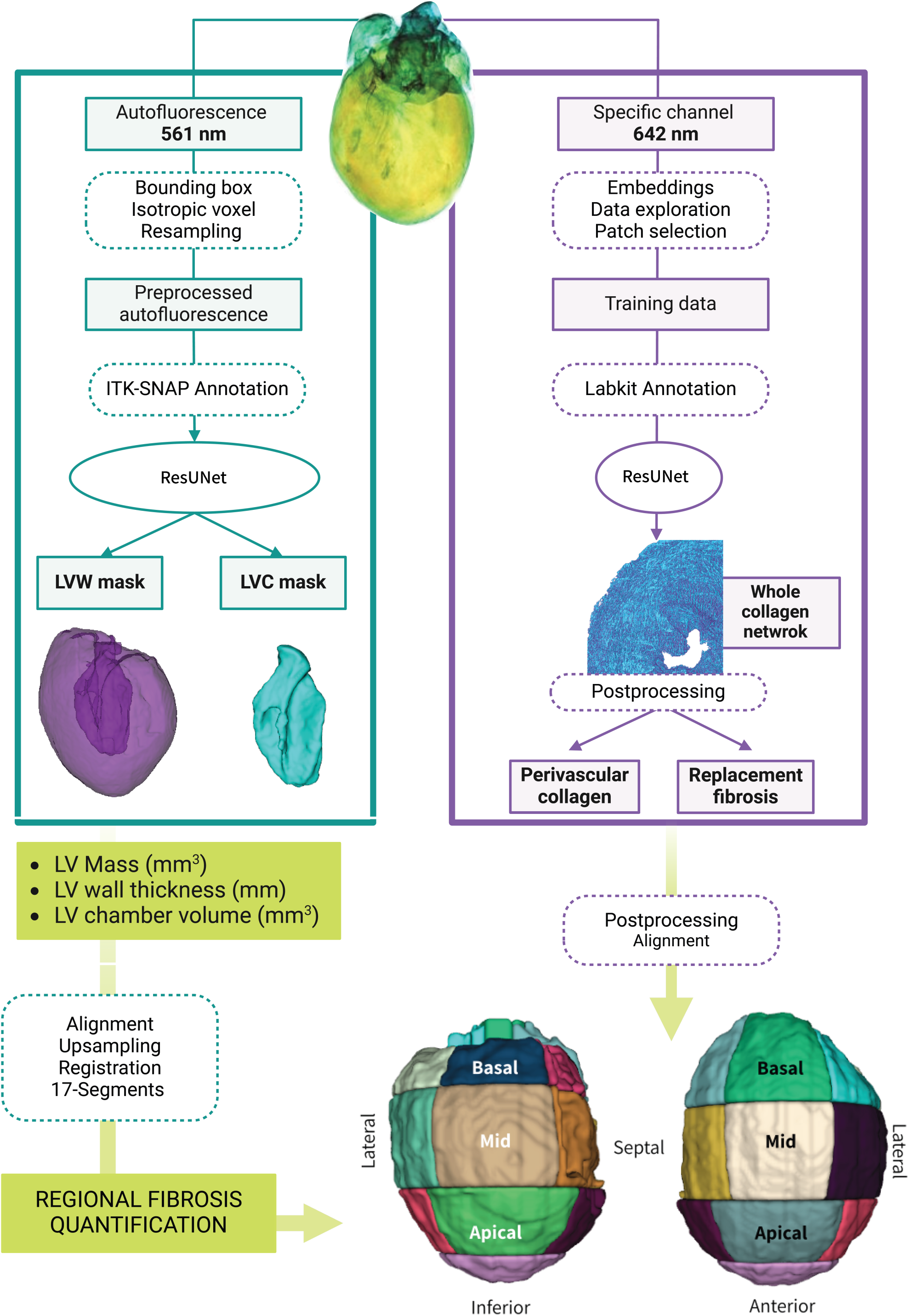
Overview of the deep learning framework for precise alignment and registration to the 17-segments of the American Heart Association for regional fibrosis quantification. (*Pipeline 1 in green on the left*) By utilizing the natural occurring autofluorescence (AF) in 561nm wavelength of whole cleared hearts (representative image at the top center) and image processing, including resampling, preprocessing and annotation for ResNet, the left ventricular wall (LVW) and chamber (LVC) are segmented. Further geometrical segmentation of the left ventricle (LV) in 17 segments of the American heart association is achieved by alignment, upsampling and registration and allows for regional fibrosis quantification. (*Pipeline 2 in purple on the right*) Collagen network segmentation based on fast green staining in 642nm wavelength of whole cleared hearts is postprocessed and enables the separation of perivascular from replacement fibrosis. Both pipelines converge to provide quantification of hypertrophy and cardiac fibrosis in regional segments of the heart, here displayed as pseudo-colored surfaces.

3D analysis of cardiac morphology was utilized to quantify the total regional LV wall thickness and chamber volume (Figure 5A-E). We found an increased LV mass in the DN/HT – vehicle group in comparison to the Healthy control group (110±16.7 mm^3^ vs 47.3±2.91 mm^3^; p<0.001), whereas semaglutide treatment led to marked reduction compared to vehicle treatment (77.8±8.84 mm^3^; p<0.001) (Figure 5F). By characterizing the volume of the 17 cardiac regions we found a uniform increase in all of the regions in the DN/HT – vehicle group in comparison to the Healthy control group, with semaglutide treatment leading to significant reduction of all of these volumes (Figure S4). Similarly, the maximum thickness of the left ventricle wall increased in the DN/HT – vehicle (1.12±0.1 mm) in comparison to the Healthy control group (0.08±0.07 mm; p<0.001), with reduction by semaglutide treatment (1.03±0.08 mm; p <0.05) (Figure 5G). This was also evident for mean LV wall thickness, which was 0.66±0.02 mm in the Healthy control group, increasing to 0.93±0.02 mm (p<0.001) in the DN/HT – vehicle group and decreasing again to 0.86 +/- 0.01 mm (p<0.05) in the DN/HT – sema group (Figure 5H). LV chamber volume in the Healthy control group was 5.9±1.07 mm^3^, increasing to 9.42±1.22 mm^3^ in the DN/HT – vehicle group (Figure 5I). Analysis in individual animals indicated large dilation of the chamber in 4 animals, with values above 15 mm^3^. Semaglutide treatment resulted in reduction (p<0.01) of the chamber volume to 4.6±0.64 mm^3^. Last, we carried out correlation analysis between LSFM scanned heart volume and heart weight recorded at termination. The results indicated strong correlation (r=0.88), thus validating that tissue processing does not alter the underlying biological differences between the groups (Figure 5J).

**Figure 5.**
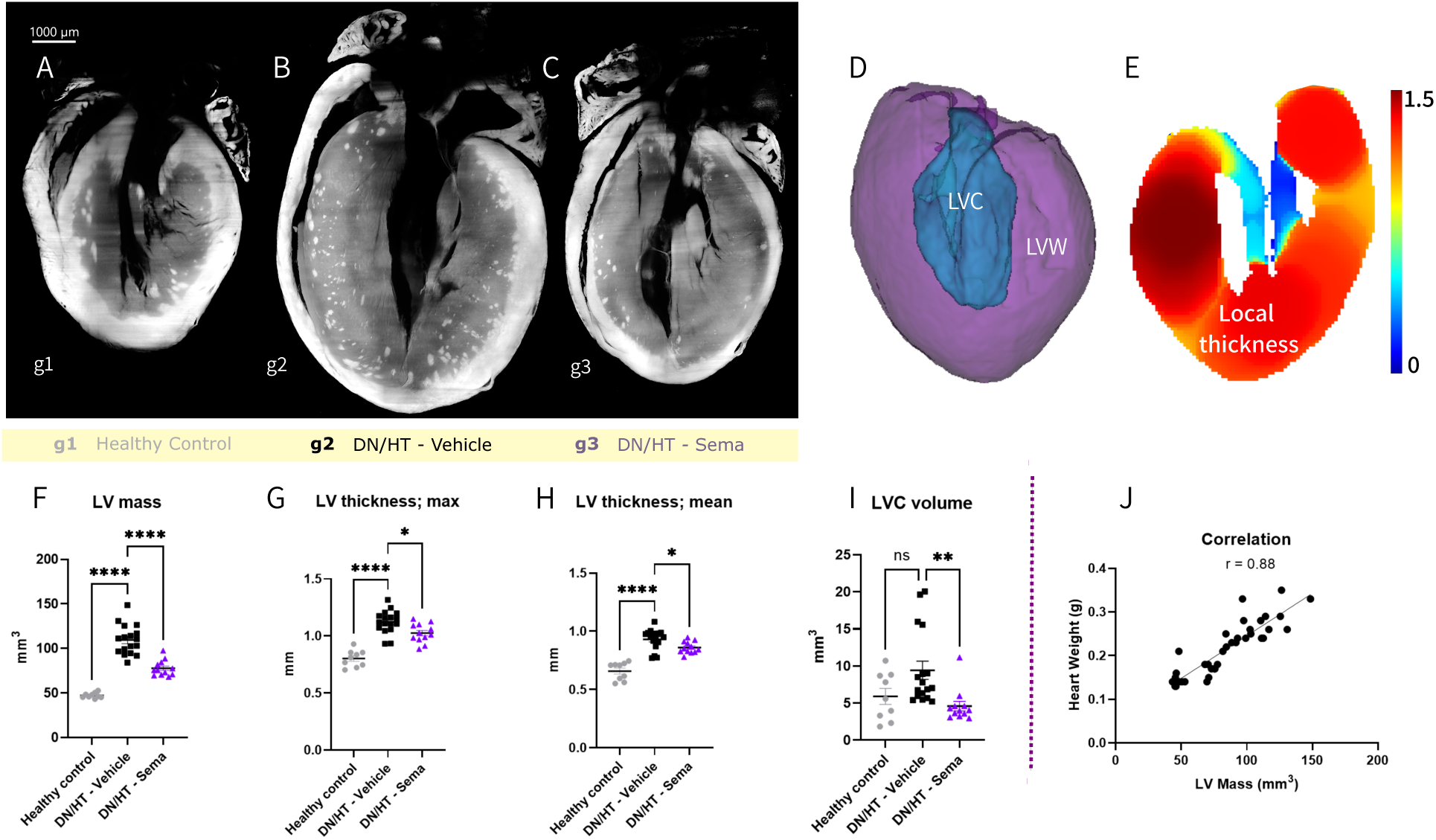
Cardiac morphometrical analysis reveals significant alterations in left ventricle (LV) metrics in a model of obesity, diabetes, and hypertrophy, further modulated by treatment with the GLP-1 antagonist semaglutide. (A-C) Representative 2D sections of hearts from (A) healthy control (*db/+)*, (B) *db/db* UNx-ReninAAV (DN/HT) *-* Vehicle treated, and (C) DN/HT – Semaglutide treated mice using LSFM whole organ imaging are displayed in grayscale of the autofluorescence (AF) channel in 561nm wavelength. (D) Representative 3D mesh projection of the left ventricle wall (LVW in purple) and left ventricle chamber (LVC in blue) segmentation derived from the deep learning framework using AF. (E) Representative image of a 2D LVW segmented volume, illustrating fitting of spheres with different radii inside the volume to quantify local thickness as a rainbow colormap with scalebar, in mm. (F-J) Cardiac morphometrical endpoints show different alterations in LV metrics, including (F) LV mass, (G) LV maximum local thickness and (H) LV mean local thickness. LVC volume (I) measurement is generated using AF segmentation and postprocessing. (J) Graph displaying a correlation between the heart weight (HW), measured at termination and the 3-dimensional LSFM imaged LV mass generated from the deep learning framework. n= 9-17, data is presented as mean +/- SEM. One-way ANOVA with Dunnet’s post-hoc test to compare means of each group to DN/HT - vehicle was applied to all data. *p<0.05; ***p<0.001; ****p<0.0001 vs Control. Graphs are generated using GraphPad Prism (v 9.2). Scale bars are 1000µm in (A-C).

The total interstitial collagen was analyzed by subtracting the perivascular collagen from the total DL detected collagen volume. There was a significant increase in interstitial collagen volume in the DN/HT –vehicle group in comparison to Healthy control (Figure 6A). The increase encompassed many cardiac segments, with the largest increase observed in mid-wall segments (Figure 6B-C, Figure S5 and S6). Treatment with semaglutide resulted in decreased collagen volume (Figure 6A, D, Figure S5 and S6). Since DN/HT model presents extensive hypertrophy we hypothesized that the increase in segmented collagen could reflect the overall expansion of the LV and thus normalized the interstitial collagen fraction to LV volume and also to individual LV segments separately. This revealed that the detected interstitial collagen in the DN/HT – vehicle group was in fact similar or lower than in the Healthy control, with no significant effect detected for semaglutide (Figure 6E-H). There is thus no increased collagen network or interstitial fibrosis in the DN/HT model compared to Healthy control, which is also supported by qualitative assessment of all the individual hearts and the corresponding DL identified segmentations (Figure 3, Figure 6I, J). The same was seen when analyzing normalized collagen content in individual cardiac regions separately (Figure S7).

**Figure 6.**
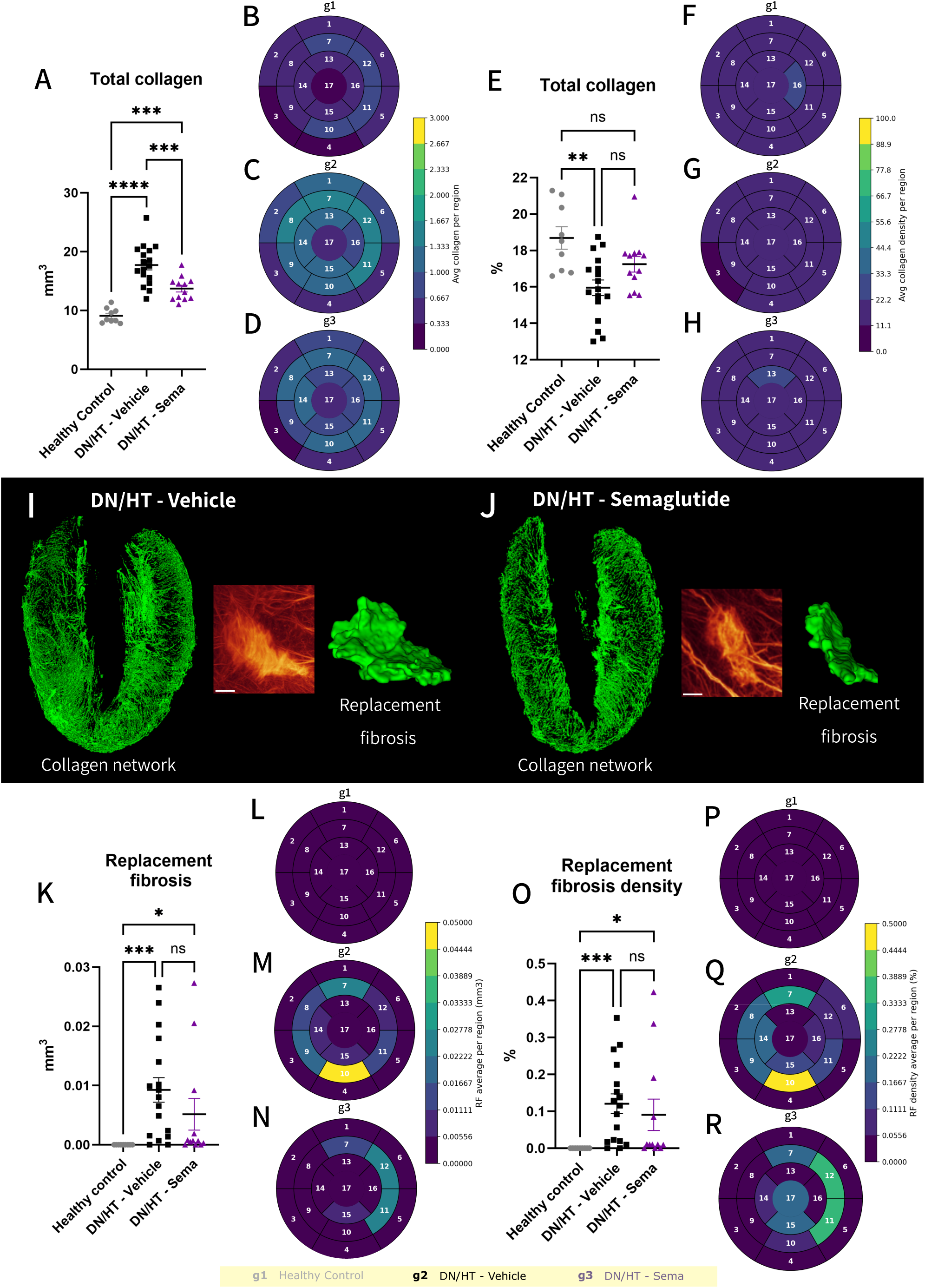
From the whole collagen network to replacement fibrosis quantification. Whole interstitial collagen network (interstitial collagen plus replacement fibrosis, excluding perivascular collagen) quantification in mm^3^ (A) and normalized to the left ventricle (LV) volume (E) together with the regional quantification average per group in 17 segments of the LV respectively (B-D; F-H). (I-J) Representative images of the interstitial collagen network segmentation in part of the LV is shown in green as maximum intensity projection and is accompanied by a representative image of the replacement fibrosis in glow colormap and its volumetric segmentation displayed as a green 3D mesh from (I) *db/db* UNx-ReninAAV (DN/HT) – Vehicle, and (J) DN/HT – Semaglutide treated mice. Replacement fibrosis quantification in mm^3^ (K) and normalized to the LV size (O) together with the regional quantification average per group in 17 segments of the LV respectively (L-N; P-R). n= 9-17, data is presented as mean +/- SEM. Kruskal-Wallis test with Dunn’s test for multiple comparisons was used for replacement fibrosis data. One-way ANOVA with Tukey’s post-hoc test to compare means of each group was applied to collagen network data. *p<0.05; ***p<0.001; ****p<0.0001 vs Control. Graphs are generated using GraphPad Prism (v 9.2). Scale bars are 150µm in I and J.

Replacement fibrosis was evident in both DN/HT – vehicle and DN/HT – sema groups, but not in the Healthy control (Figure 3, Figure 6I-J, Figure S10). At the location of replacement fibrosis, the cardiomyocytes that were distinguishable in autofluorescence wavelength were lost (Figure S8). Quantification of total replacement fibrosis volume in the entire LV indicated no significant effect for semaglutide, largely due to higher values observed in four animals (Figure 6K-N). The same was seen when replacement fibrosis volume was normalized to the total LV volume (Figure 6O-R). Both DN/HT – vehicle and DN/HT – sema differed significantly from Healthy control group in total and normalized replacement fibrosis volume. Data for individual cardiac regions indicated large variability in where replacement fibrosis develops, with no clear pattern detected (Figure S9).

Last, we studied perivascular fibrosis. We found a significant increase in perivascular collagen volume in the DN/HT – vehicle group in comparison to Healthy control (Figure 7A-C). Perivascular collagen content was highest in the anterior and anterolateral cardiac regions (Figure 7B-C; Figure S10). Importantly, semaglutide resulted in a marked reduction in perivascular fibrosis in comparison to DN/HT – vehicle (Figure 7A-D). To account for possible expansion in vascular length in cardiac hypertrophy, we also normalized the perivascular collagen volume to vascular length, which further supported the increased perivascular collagen in the DN/HT – vehicle group in comparison to Healthy control and demonstrated that semaglutide treatment led to reduction of perivascular fibrosis to the level of Healthy control (Figure 7E-H).

**Figure 7.**
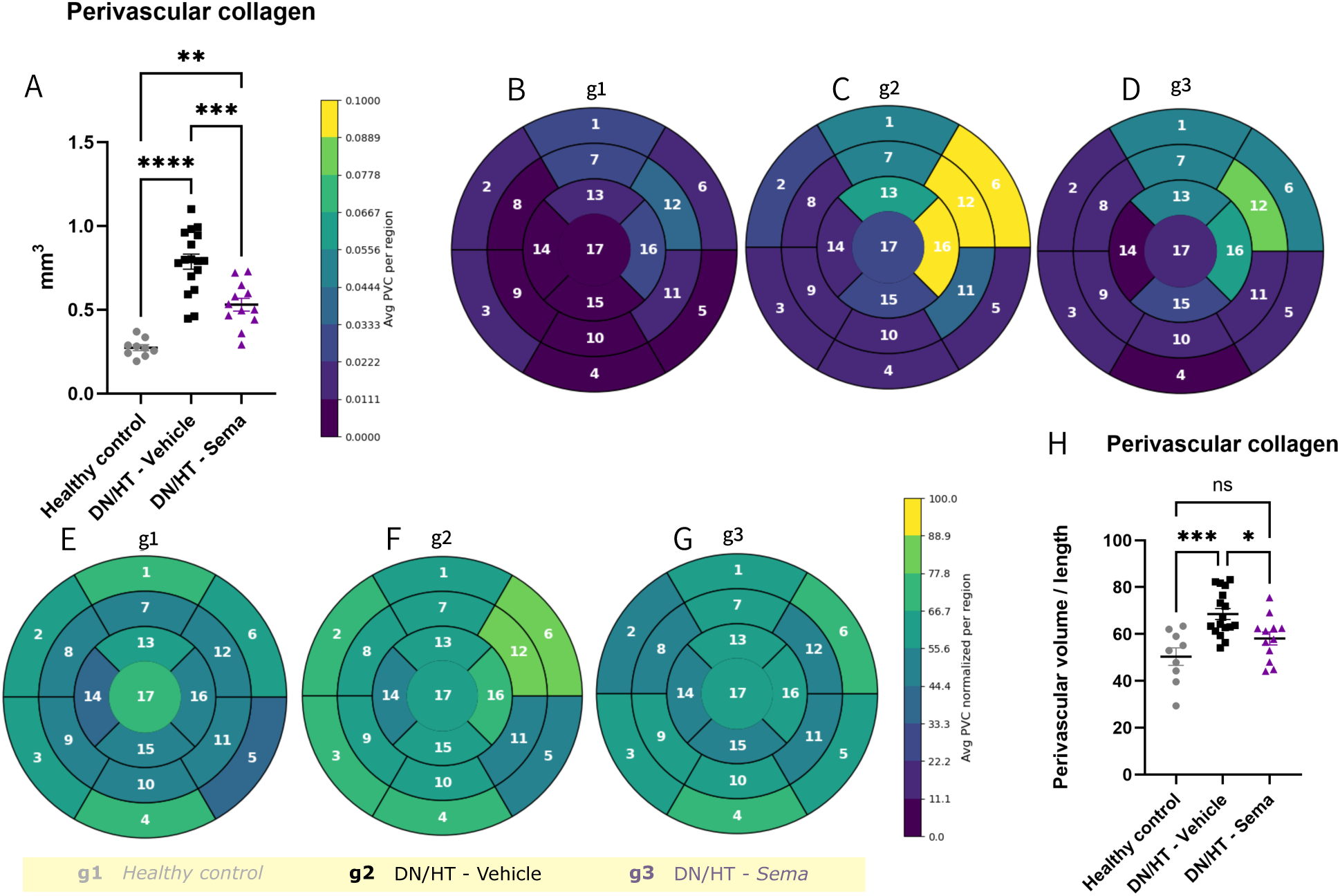
Semaglutide treatment reduces perivascular collagen fibrosis. Graphs display the total perivascular collagen quantification in mm^3^ (A) and normalized to the vessel length (H) in Healthy controls (*db/+)* and *db/db* UNx-ReninAAV (DN/HT) animals treated with vehicle or semaglutide. n= 9-17, data is presented as mean +/- SEM. Regional group averages of the quantification in (A) and (H) in 17 segments of the LV respectively (B-G). One-way ANOVA with Tukey’s post-hoc test to compare means of each group. *p<0.05; ***p<0.001; ****p<0.0001 vs Control. Graphs are generated using GraphPad Prism (v 9.2).

## Discussion

We have demonstrated here a 3D LSFM and AI-based quantitative platform for rapid assessment of fibrosis in entire mouse hearts. This method overcomes the resolution limits of preclinical *in vivo* imaging modalities and the selection bias of histology methods arising from the sectioning of the heart. It enables distinguishing between perivascular, interstitial and replacement fibrosis. We validated the methods on a mouse model with diabetes, renal failure and hypertension, detecting prevalent cardiac hypertrophy and perivascular fibrosis. To validate the sensitivity of the method to detect the efficacy of pharmaceuticals we applied it to characterize the effect of chronic semaglutide treatment. We demonstrated a significant reduction of cardiac hypertrophy and perivascular fibrosis in semaglutide treated group, whereas instances of replacement fibrosis could still be detected in the compound treated group.

Laboratory mice represent the model of choice in preclinical drug discovery research, including in cardiovascular diseases. Yet, in comparison to the human heart, the small size of mouse heart (diameter less than 6 mm) and high heart rate (500-700 beats per minute) pose unique challenges for imaging methods that are commonly applied in clinical practice. In particular, accurate quantitative assessment of cardiac fibrosis has been difficult due to the sensitivity and resolution limits of *in vivo* imaging modalities that need a high image acquisition rate to overcome the movement of the heart. For this reason, attention has been put on collagen- and elastin-specific molecular imaging agents for use in cardiac magnetic resonance imaging (CMR) and positron emission tomography (PET) and which have helped in identifying large fibrotic scars, developing in myocardial infarction (Helm et al., 2008; Wildgruber et al., 2014). However, the resolution has remained insufficient to distinguish interstitial and perivascular fibrosis developing in HFpEF models and neither of these methods have been validated in pharmacological research. Histology on tissue sections has remained the method of choice until now but suffers from several challenges (Schipke et al., 2017). The random sampling of sections for stereological charting of cardiac fibrosis is often not suitable as the fibrosis is not uniformly distributed in the myocardium, whereas analysis of specific planes may not capture the stochastically developing fibrosis (Schipke et al., 2017). Finding more sensitive means to image and quantify MF has thus been defined as a key priority in heart failure research (de Boer et al., 2019) . The resolution achieved with the imaging method described here surpasses what is currently possible with *in vivo* imaging and compared to histology, enables the reconstruction of the 3D arrangement of collagen distribution in the heart and captures stochastic fibrotic events that may remain undetected in tissue sections. In the healthy heart, our results support the knowledge from scanning electron microscopy (Kanzaki et al., 2010). We could visualize supportive collagen network surrounding cardiomyocytes and blood vessels. 3D imaging of the pericardium illustrated a multi-directional network of collagen in detail that has remained difficult if not impossible to achieve before.

AI is revolutionizing biomedical research and clinical practice and in the context of cardiovascular diseases and CMR was found to be equal or superior to experts in diagnostics (Wang et al., 2024). Progress has been made in applying AI to detect fibrosis in clinical imaging data (Penso et al., 2023; Wang et al., 2024). DL methods are gaining popularity in quantifying fibrosis on tissue sections and have been applied to characterize FG staining in a mouse heart failure model, showing strong correlation with collagen 1a1 protein content detected in Western Blot (Remes et al., 2023). Here we implemented DL to identify collagen content in the 3D imaged heart at high resolution and detect both replacement and perivascular fibrosis, in addition to the assessment of total collagen content in the tissue. Furthermore, we used AI in automated analysis of cardiac hypertrophy and LV chamber dilation. To enhance clinical translatability, we divided the left ventricle into 17 segments. Overall, the developed sample preparation, imaging and image analysis pipeline is suitable for high-throughput workflow. The samples can be batch processed and require only simple liquid change steps that are possible to implement on liquid handling robots. Whole heart imaging of FG can be performed in less than 30 minutes and image analysis requires minimal operator input. The overall time thus compares favorably to standard histological assessment of fibrosis.

We show that the DN/HT mouse model that has been characterized for hypertension, renal failure and diabetes endpoints (Harlan et al., 2018; Østergaard et al., 2021b; Dalbøge et al., 2022) also develops hypertrophy and MF. As the model carries the common HFpEF co-morbidities (obesity, diabetes mellitus type 2, hypertension, kidney disease) and progresses over a long period (12 weeks), it is well suited to study the mechanisms of chronic HFpEF. We found the mouse model to develop extensive perivascular fibrosis, in addition to replacement fibrosis. Perivascular fibrosis is recognized in biopsies and post-mortem analysis in hypertensive heart disease, hypertrophic heart disease, diabetes and HFpEF (van Hoeven and Factor, 1990; Shimizu et al., 1993; Dai et al., 2012). Coexistence of diabetes and hypertension has been found to increase fibrosis burden in comparison to having either diabetes or hypertension alone (van Hoeven and Factor, 1990). The development of extensive perivascular fibrosis has been hypothesized to contribute to tissue hypoxia, resulting in localized ischemia, myocyte necrosis and replacement fibrosis (van Hoeven and Factor, 1990; Shimizu et al., 1993; Kwong et al., 2008; de Boer et al., 2019). Focal replacement fibrosis is also common in HFpEF CMR findings, together with localized ischemic events (Kato et al., 2015b; Kanagala et al., 2019). Qualitative assessment of 3D imaged hearts supports this hypothesis as the replacement fibrosis was often in contact with or adjacent to perivascular fibrosis areas. Our data does not support increased density of interstitial collagen fibrils in the DN/HT mouse model in comparison to the Healthy control group as the total increase in collagen content per region is explained by cardiac hypertrophy, rather than increase in collagen density. Region-wise analysis indicated that all 17 cardiac regions increase in volume in the DN/HT mouse model in comparison to the Healthy control. While all regions also demonstrated an increase in total collagen content in comparison to the Healthy control, then the increase was abolished after normalization to region volume. Interestingly, characterization of fibrosis in diabetic *db/db* mice fed high fat diet, developing HFpEF symptoms but without high hypertension and kidney disease, revealed primarily perivascular fibrosis. The study did not detect replacement fibrosis or extensive collagen accumulation in the interstitium (Alex et al., 2018). These results suggest that the renal and hypertension components promote the progression of fibrosis beyond the cardiac vasculature. Mainly perivascular fibrosis has also been found in models of obesity (Zaman et al., 2004; Fopiano et al., 2021).

Most preclinical studies in mice, including research in HFpEF models, have not distinguished between the different fibrosis subtypes or have only separated perivascular fibrosis. The reason behind this is the inherent difficulty in conclusively determining whether the staining that is evident on a tissue section represents an edge of a replacement fibrosis scar, a single interstitial fibrosis area or even a border of a vascular area from the neighboring tissue plane. Thus, the here developed 3D imaging method can help to improve the sensitivity in MF analysis to identify the efficacy of tested drugs to reduce the different MF subtypes.

GLP-1R agonists show benefits in clinical HF trials but their mode of action in the heart requires further characterization. Here we found that chronic semaglutide administration reduces perivascular fibrosis in HFpEF. Reduction of replacement fibrosis did not reach significance as focal fibrosis was still detectable in the semaglutide treatment arm. Replacement fibrosis is generally considered irreversible. Temporal CMR analysis of replacement fibrosis in dilated cardiomyopathy patients under optimal treatment regimen showed stable replacement fibrosis over 14-month period (Rubiś et al., 2024). The results from our data suggest that initial fibrotic areas likely developed in the model initiation period and did not respond to semaglutide. A number of previous studies have assessed GLP-1R agonists in rodent HF models. Liraglutide was found to reduce cardiac hypertrophy and total fibrosis content in a HFpEF mouse model infused with AngII and fed high-fat diet (Gaspari et al., 2016; Withaar et al., 2021). Similarly, in a hypertension model of mice infused with AngII alone, liraglutide reduced collagen accumulation (Chen et al., 2021). Positive effects for liraglutide on cardiac fibrosis and hypertrophy were found in a mouse transverse aortic constriction model of pressure overload induced HF (Li et al., 2024). Likewise, semaglutide reduced cardiac fibrosis and hypertrophy, in line with broad cardiometabolic benefits and extensive changes in gene expression (Withaar et al., 2023; Ma et al., 2024). Our data expands this knowledge to mouse model of diabetes, obesity, hypertension and chronic kidney disease associated with HF and demonstrates benefits on reducing cardiac hypertrophy and perivascular fibrosis.

While our study shows a number of benefits for 3D fibrosis imaging and DL-assisted analysis and indicates beneficial anti-fibrotic effects for semaglutide, it also comes with limitations that we recognize. First, FG staining, although highly specific to collagen (Timin and Milinkovitch, 2023), does not distinguish the different types of collagen protein. The composition of extracellular matrix is known to change in the progression of HF, which the described method here remains insensitive towards (Li et al., 2018). At the same time, antibodies targeting abundant extracellular matrix proteins, like collagen, fail to provide uniform staining in large and dense cleared tissue samples. We also analyzed a single time-point for MF and lack evidence on temporal changes and how MF measurements correlate with changes in cardiac function.

In conclusion, we have demonstrated here a 3D imaging method for rapid assessment of collagen content in entire rodent hearts for HF research. We provide DL-based image analysis workflow to quantify total, perivascular and replacement cardiac fibrosis in different heart segments and in parallel assess hypertrophy. We show the beneficial effect of semaglutide in reducing perivascular fibrosis and cardiac hypertrophy in a mouse model with diabetes, renal failure and hypertension associated HF. We propose that the developed methods will simplify and accelerate basic and drug discovery research for heart diseases and could also be implemented to characterize fibrosis in other diseases.

## Acknowledgements

The authors would like to thank Neringa Zitkute and Sabrina Krarup for assisting with sample preparation.

## Author contributions

Investigation, Methodology: S.B.B., S.T.Y., M.H., M.B.R., D.M.J., M.C., L.S.T. Data curation: S.B.B., C.G.S. Visualization: S.B.B., S.T.Y., M.H. Supervision: H.L.H., C.G.S., T.B.S, U.R. Writing – original draft: S.B.B., U.R., Writing – review & editing: S.B.B., S.T.Y., M.H., M.B.R., D.M.J., M.C., L.S.T., H.L.H., G.T., T.B.S., C.G.S. and U.R. Project administration: M.C., G.T., H.L.H., C.G.S., U.R. Conceptualization: U.R.

