## Supplementary figures for "Deep Learning-Enhanced 3D Imaging Unveils Semaglutide Impact on Cardiac Fibrosis"

Figure S1

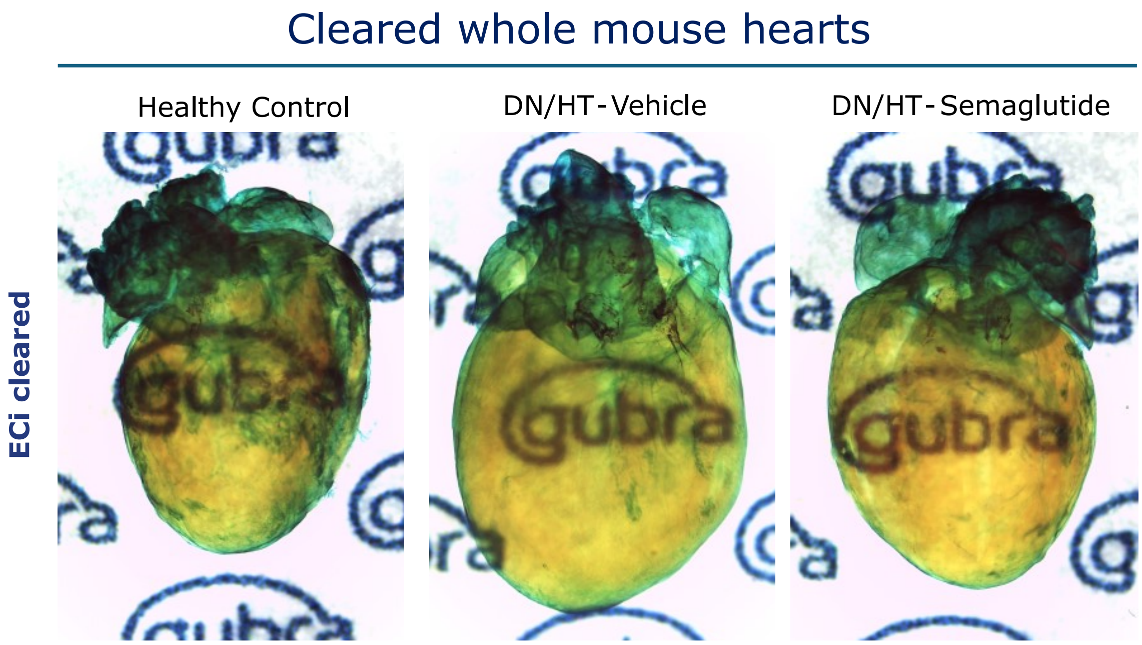

Cleared whole mouse hearts
Healthy Control
DN/HT-Vehicle
DN/HT-Semaglutide
ECi cleared

Figure S2

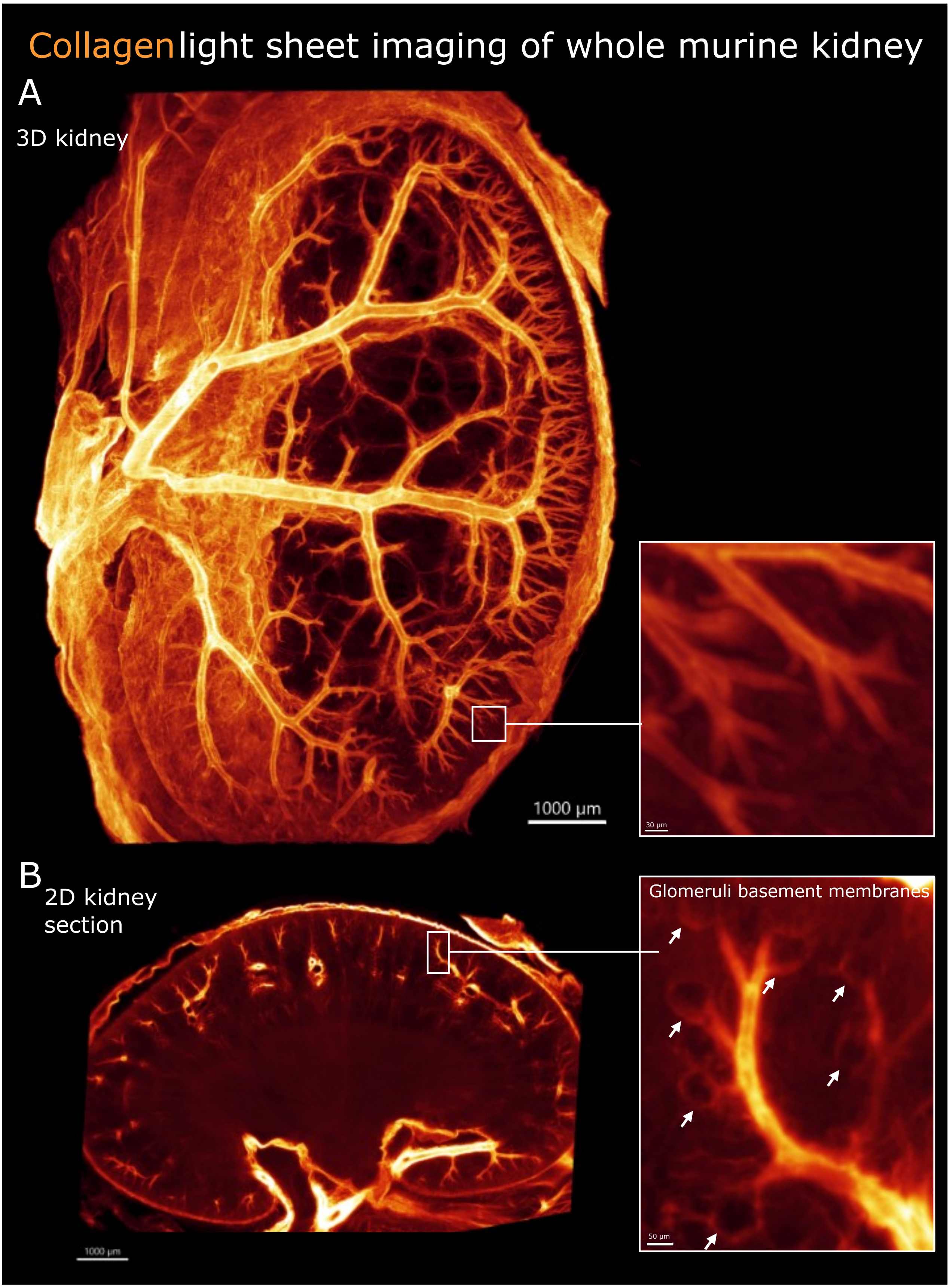

Collagen light sheet imaging of whole murine kidney
A
3D kidney
1000 µm
30 µm
B
2D kidney section
1000 µm
Glomeruli basement membranes
50 µm

Figure S3

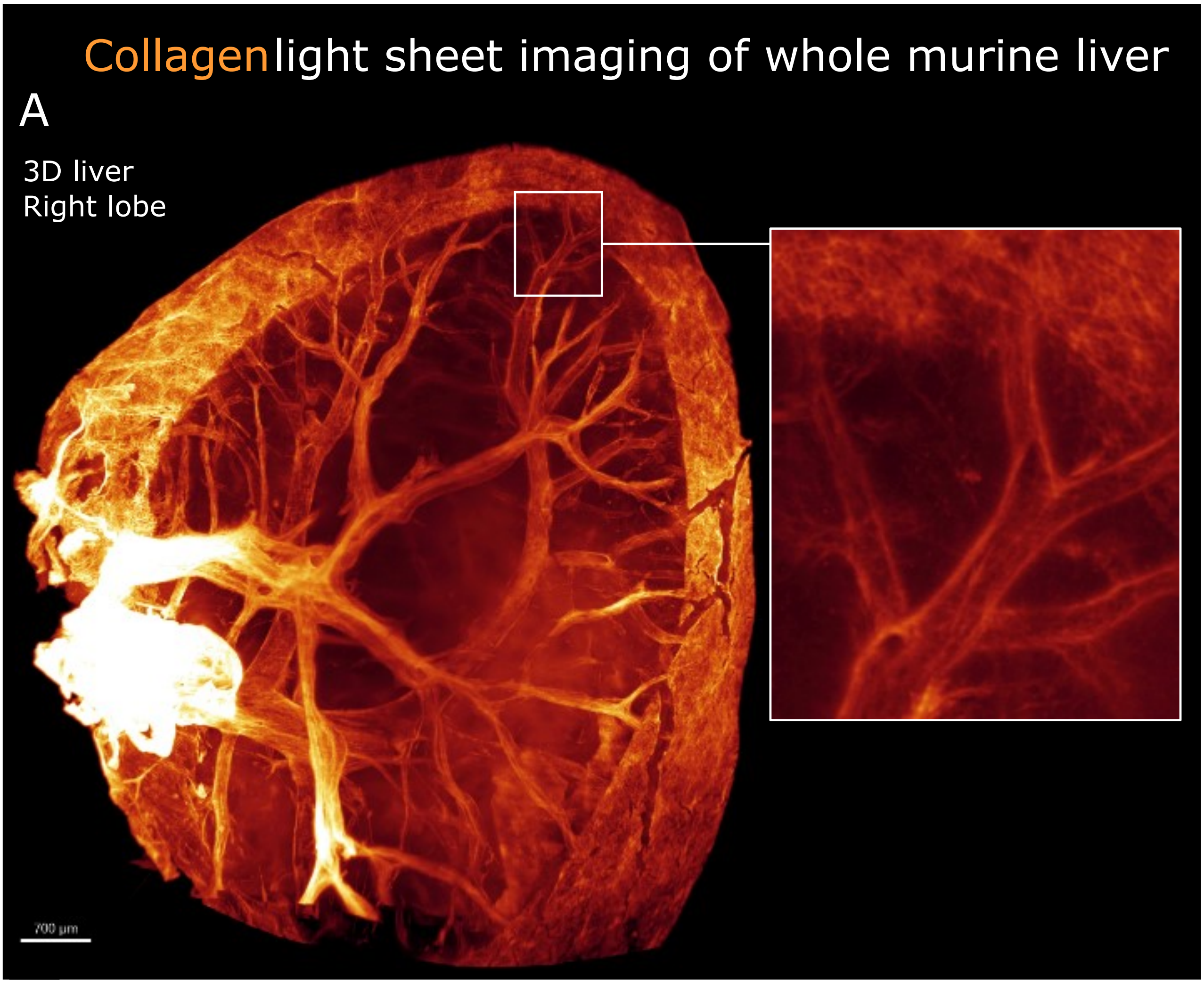
Collagen light sheet imaging of whole murine liver
A
3D liver
Right lobe
700 µm

Figure S4

### Regional hypertrophy

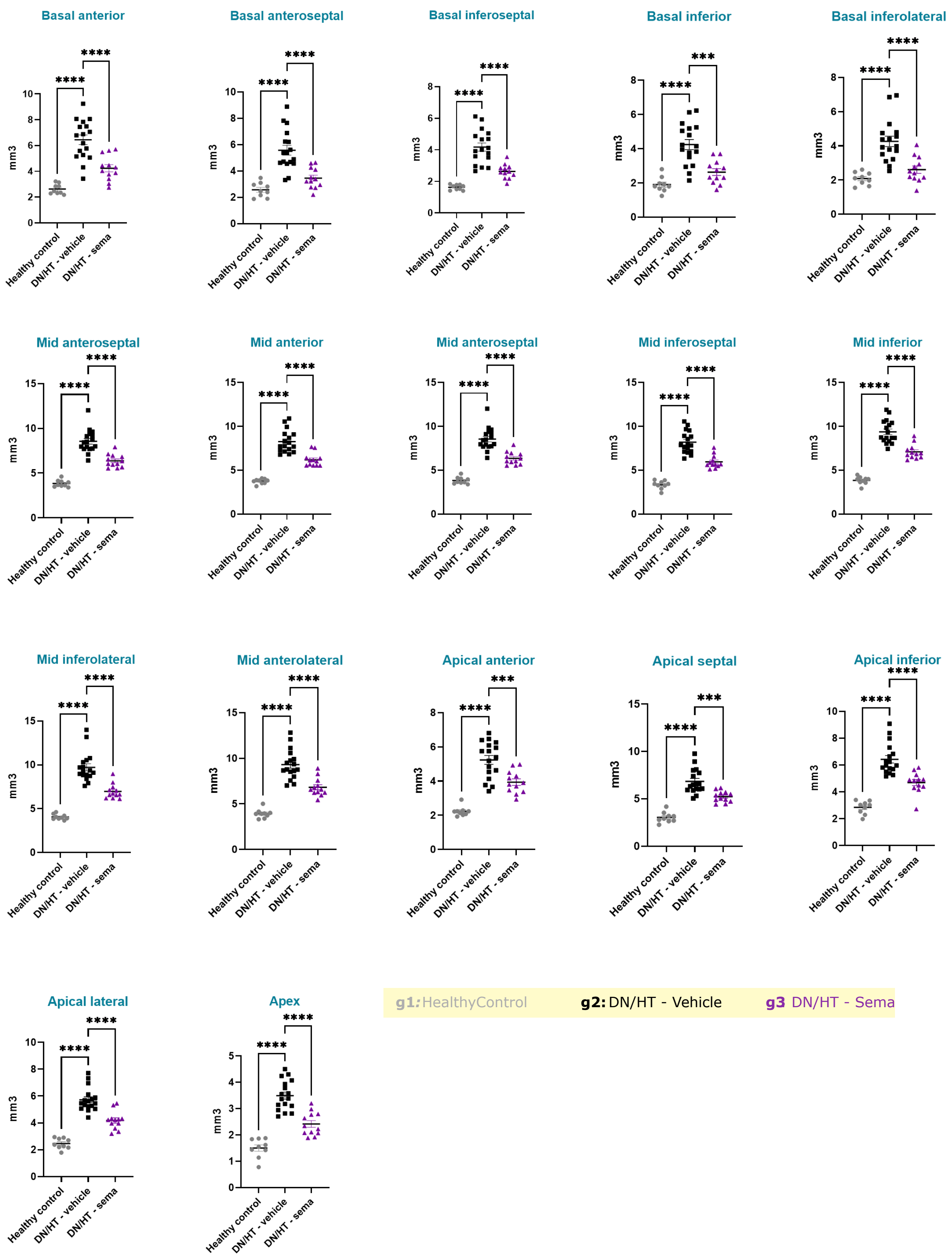

Figure S5

### Regional collagen

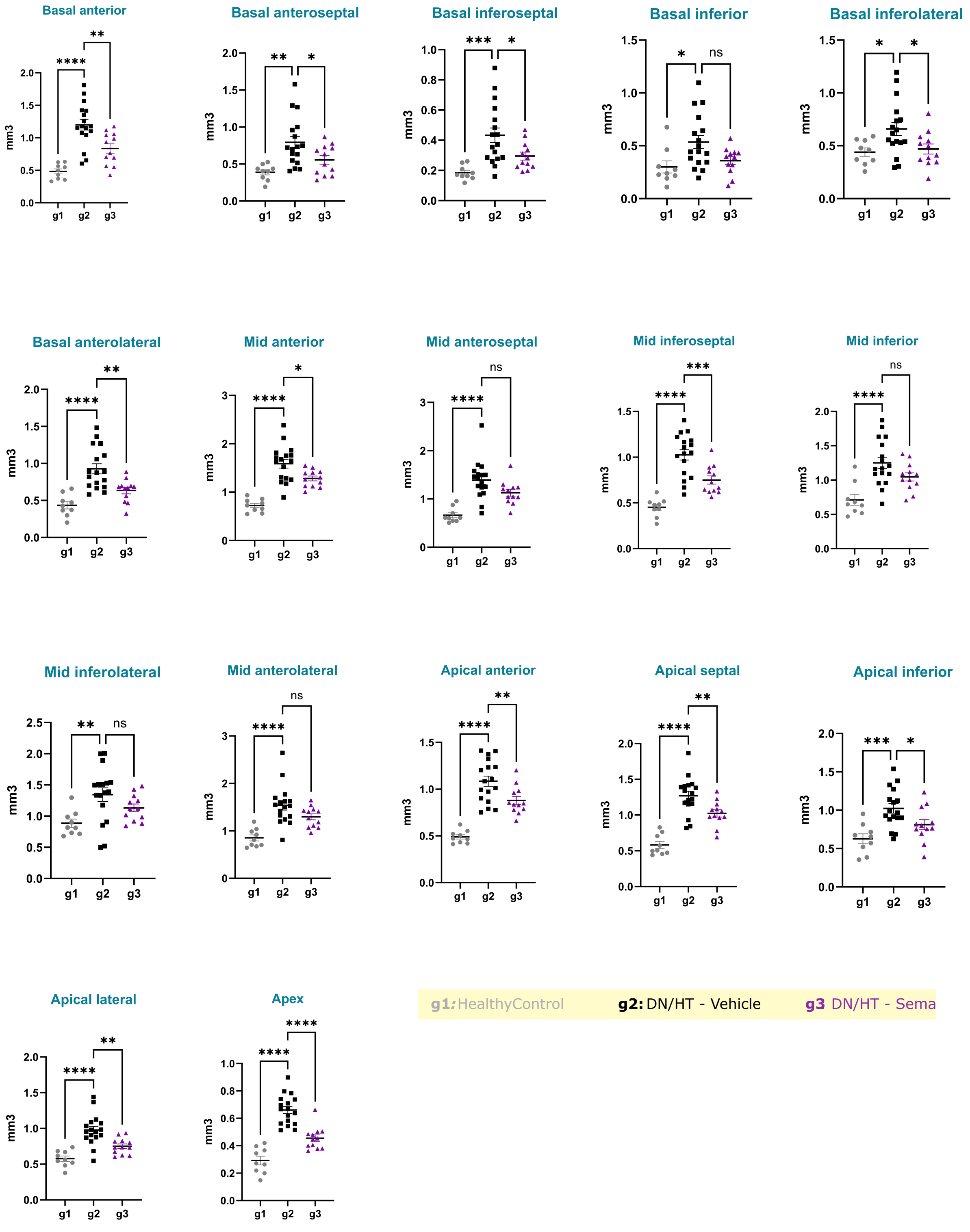

Figure S6

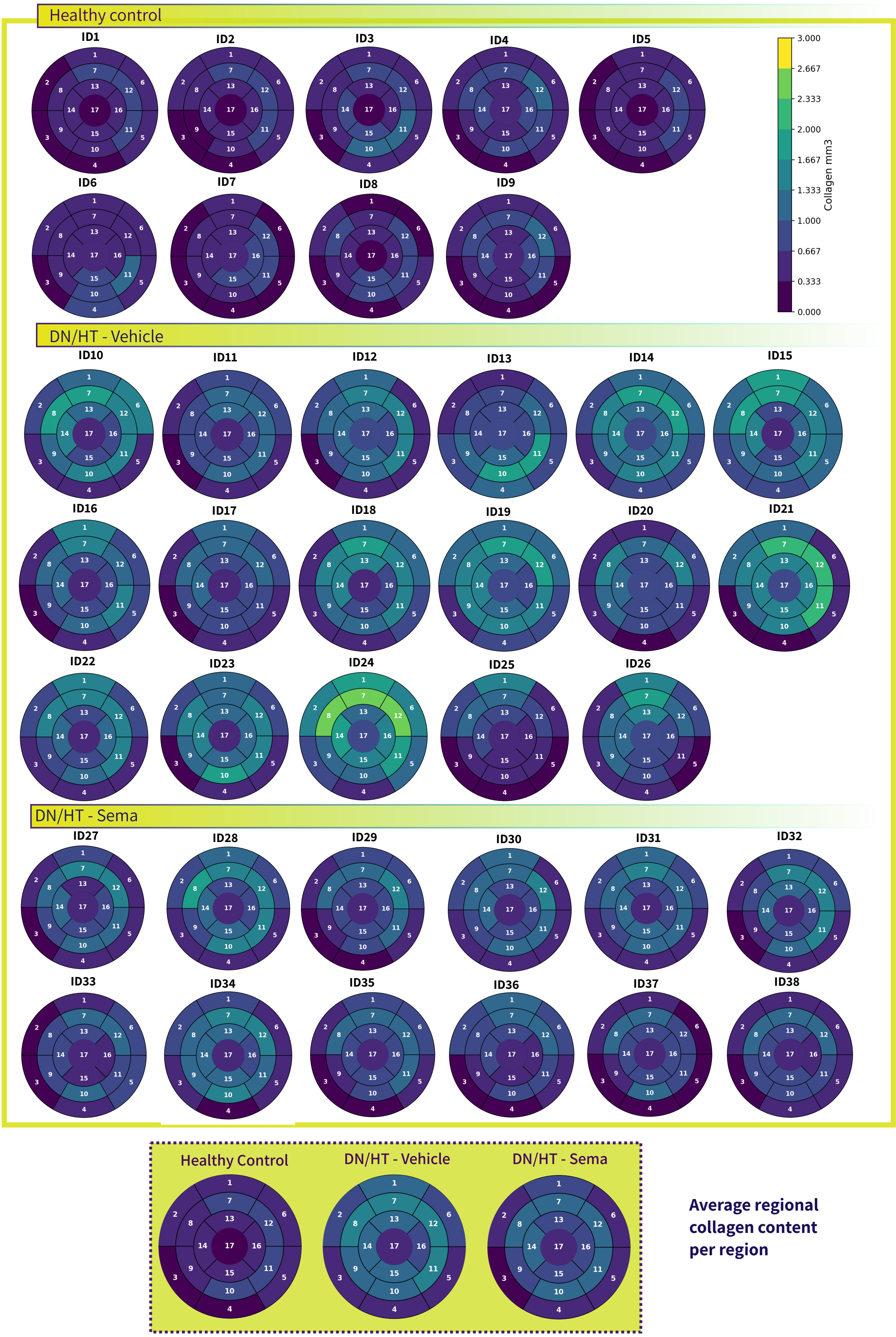
Healthy control
ID1
ID2
ID3
ID4
ID5
ID6
ID7
ID8
ID9
3.000
2.667
2.333
2.000
1.667
1.333
1.000
0.667
0.333
0.000
Collagen mm3
DN/HT - Vehicle
ID10
ID11
ID12
ID13
ID14
ID15
ID16
ID17
ID18
ID19
ID20
ID21
ID22
ID23
ID24
ID25
ID26
DN/HT - Sema
ID27
ID28
ID29
ID30
ID31
ID32
ID33
ID34
ID35
ID36
ID37
ID38
Healthy Control
DN/HT - Vehicle
DN/HT - Sema
Average regional collagen content per region

Figure S7

### Regional collagen density

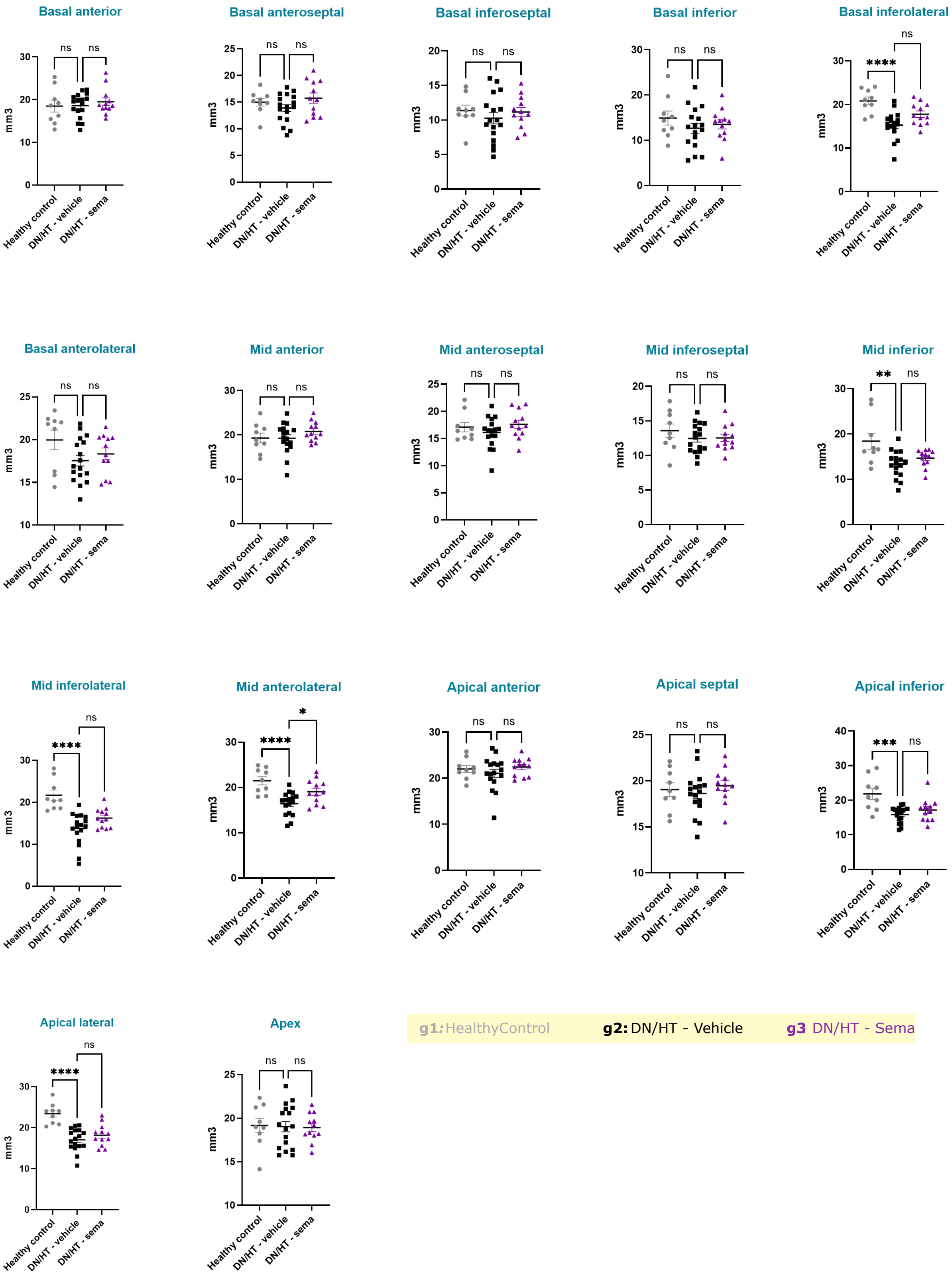

Figure S8

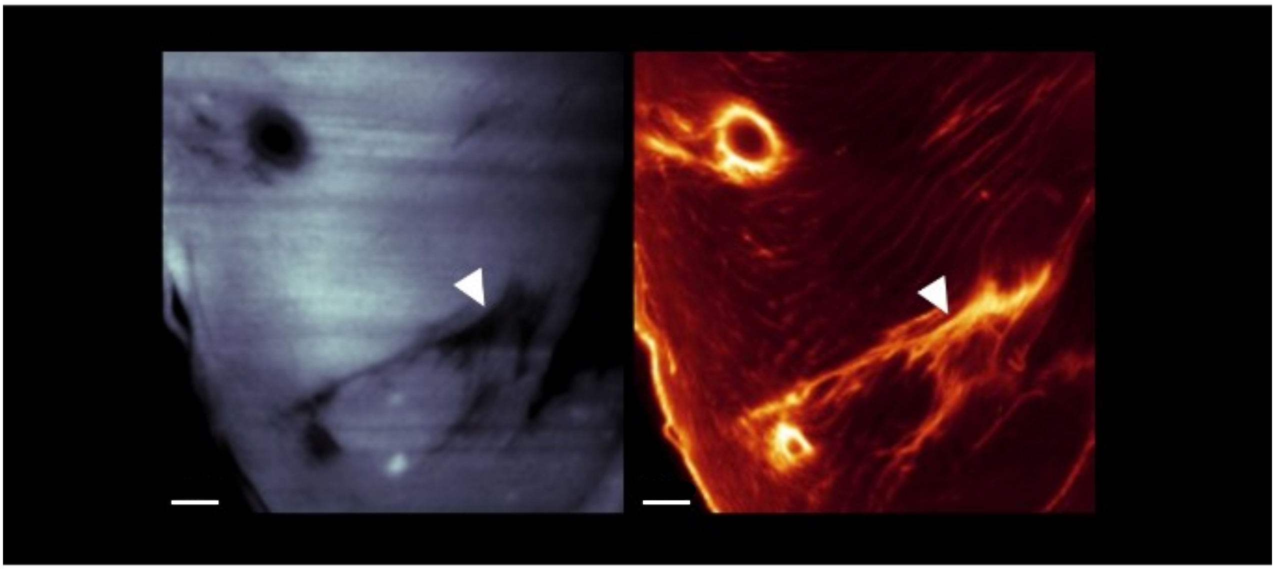

Figure S9

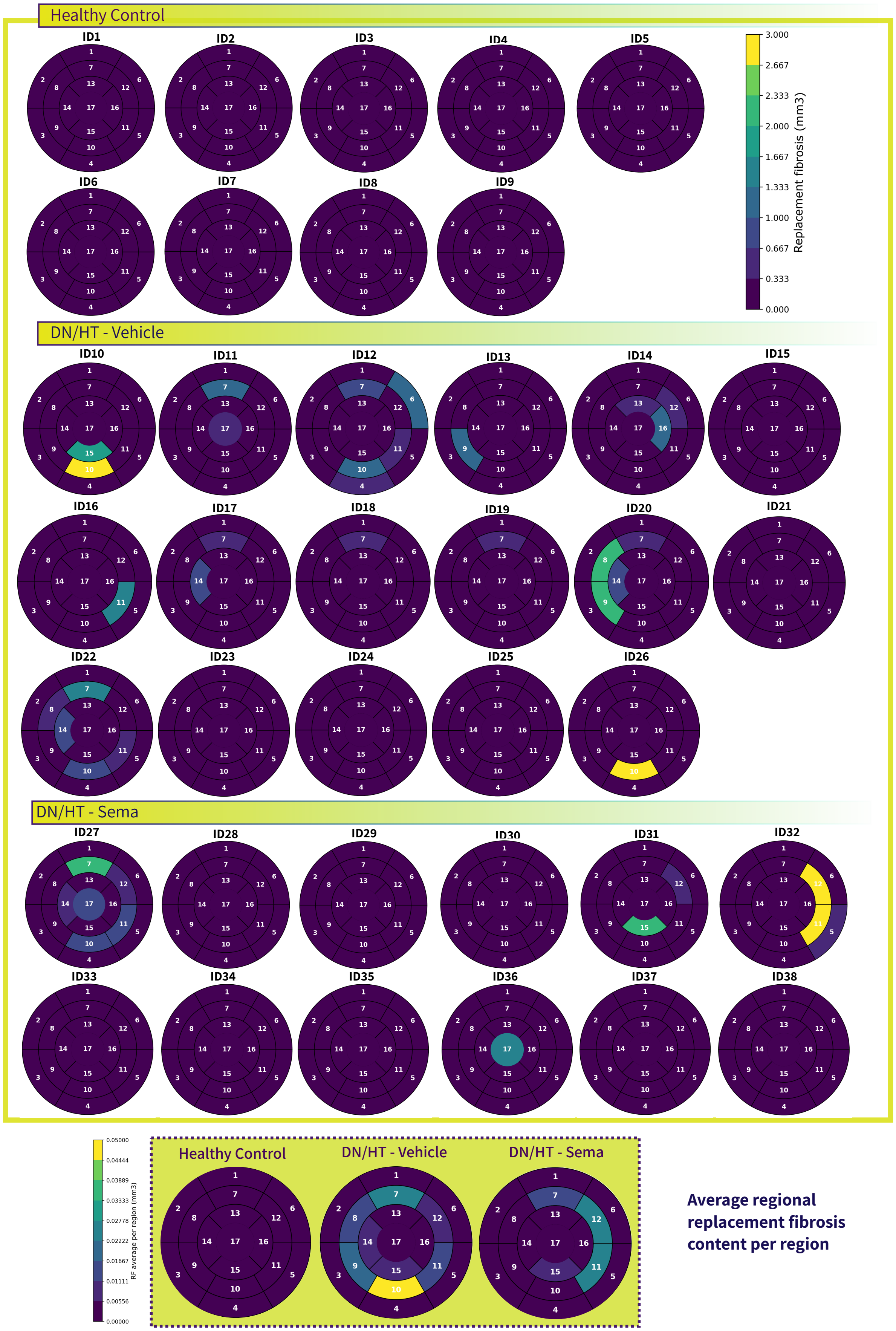

Healthy Control
ID1
ID2
ID3
ID4
ID5
ID6
ID7
ID8
ID9
3.000
2.667
2.333
2.000
1.667
1.333
1.000
0.667
0.333
0.000
Replacement fibrosis (mm3)
DN/HT - Vehicle
ID10
ID11
ID12
ID13
ID14
ID15
ID16
ID17
ID18
ID19
ID20
ID21
ID22
ID23
ID24
ID25
ID26
DN/HT - Sema
ID27
ID28
ID29
ID30
ID31
ID32
ID33
ID34
ID35
ID36
ID37
ID38
0.05000
0.04444
0.03889
0.03333
0.02778
0.02222
0.01667
0.01111
0.00556
0.00000
RF average per region (mm3)
Healthy Control
DN/HT - Vehicle
DN/HT - Sema
Average regional replacement fibrosis content per region

Figure S10

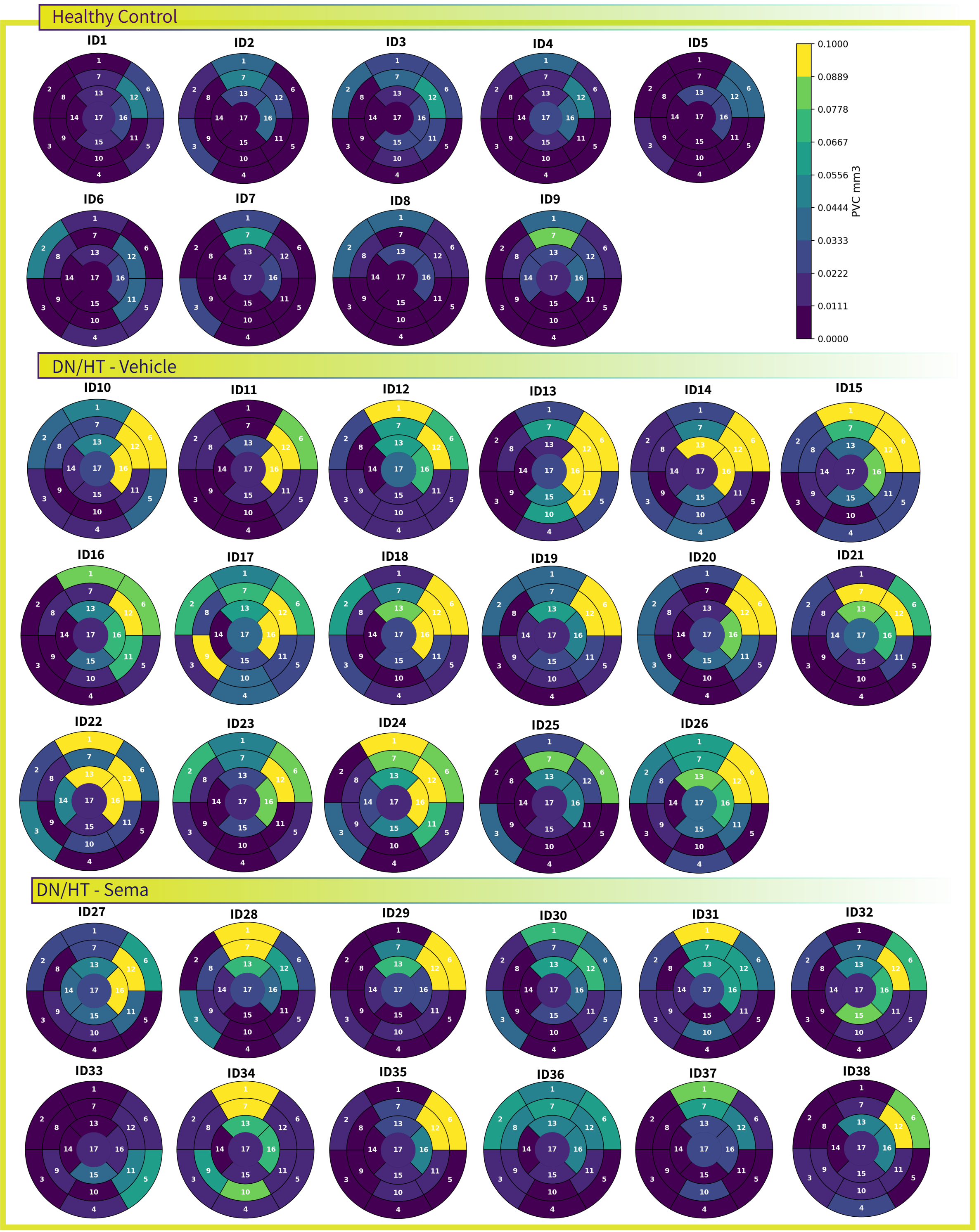


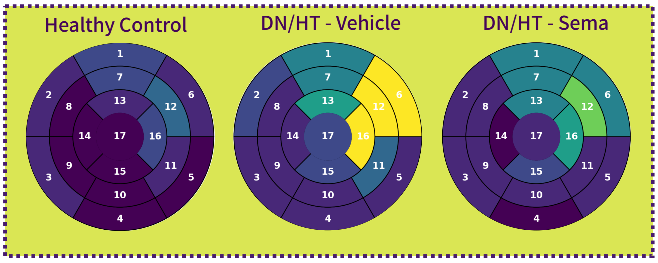


**Average regional perivascular collagen content per region**
