## Supplementary figure legends for "Deep Learning-Enhanced 3D Imaging Unveils Semaglutide Impact on Cardiac Fibrosis"

**Supplemental Figure Legends**

**Figure S1. Refractive index (RI) matching whole mouse hearts.** Optically cleared whole mouse hearts with ethyl cinnamate (ECi) to match the refractive index of fixed proteins and lipids allowing for high-resolution 3D imaging of collagen in intact tissue.

**Figure S2. 3D light sheet fluorescence microscopy in mouse kidney with Fast Green (FG) collagen staining.** Representative 3D light sheet fluorescence microscopy (LSFM) images of collagen distribution in a healthy C57BL6/J mouse kidney, labeled using FG in glow. (A) A maximum intensity projection (MIP) of a 1 mm thick slice through the 3D imaged kidney in long-axis plane, with zoomed vasculature on the right. (B) 2D cross-sectional image of a kidney with a magnified view to display glomerular basement membrane labelling (arrows). Scale bars are 1000μm (A, B) and 100μm in boxed images.

**Figure S3. 3D light sheet fluorescence microscopy in mouse liver with Fast Green (FG) collagen staining.** Representative 3D light sheet fluorescence microscopy (LSFM) images of collagen distribution in a healthy C57BL6/J mouse liver, labeled using FG in glow. (A) A maximum intensity projection (MIP) of a 1 mm thick slice through the 3D imaged liver in long-axis plane, shows the vasculature with FG highlighting collagen. Scale bar, 700μm.

**Figure S4. Regional hypertrophy.** Region size in $mm^3$ of each segment in the left ventricle (LV)**.** One-way ANOVA with Dunnet's post-hoc test to compare means of each group to the *db/db* UNxReninAAV Vehicle was applied to all data. *p<0.05; ***p<0.001; ****p<0.0001. Graphs are generated using GraphPad Prism (v 9.2).

**Figure S5. Regional collagen**. Total interstitial collagen content per region in the 17 left ventricle (LV) segments. One-way ANOVA with Dunnet's post-hoc test to compare means of each group to the Vehicle was applied to all data. *p<0.05; ***p<0.001; ****p<0.0001 vs *db/db* UNxReninAAV Vehicle. Graphs are generated using GraphPad Prism (v 9.2).

**Figure S6**. **Regional collagen content.** Total interstitial collagen content (interstitial and replacement fibrosis) mapped to cardiac segments in individual animals from Healthy control (db/+), *db/db* UNx-ReninAAV (DN/HT) – Vehicle, and *DN/HT* – Semaglutide groups.

**Figure S7**. **Regional collagen density.** Total interstitial collagen content per region in the 17 left ventricle (LV) segments normalized to region volume. One-way ANOVA with Dunnet's post-hoc test to compare means of each group to the *db/db* UNxReninAAV (DN/HT) – Vehicle was applied to all data. *p<0.05; ***p<0.001; ****p<0.0001. Graphs are generated using GraphPad Prism (v 9.2).

**Figure S8. Replacement fibrosis characterization**. 2D section overview (from 3D imaged heart), illustrating replacement fibrosis in the myocardium. (Left) Autofluorescence with arrowhead depicting the absence of cardiomyocytes. (Right) Collagen staining with FG with arrowhead showing replacement fibrosis. Scale bars: 300 μm.

**Figure S9. Segment plots for replacement fibrosis.** Replacement fibrosis mapped to cardiac segments in individual animals from Healthy control (*db/+*)*, db/db* UNx-ReninAAV (DN/HT) – Vehicle, and (DN/HT) – Semaglutide groups.

**Figure S10. Segment plots for all animals for perivascular fibrosis.** Perivascular fibrosis mapped to cardiac segments in individual animals from Healthy control (*db/+*)*, db/db* UNx-ReninAAV (DN/HT) – Vehicle, and DN/HT – Semaglutide groups.
